# Cross-species chromatin accessibility pinpoints HIVEP3 as a driver of thymic epithelial maturation and self-antigen expression, preventing chronic inflammation

**DOI:** 10.64898/2026.06.16.732438

**Authors:** Sümeyye Yayilkan, Marie Massoda, Séverine Ménoret, Axel You, Lucas Brusselle, Claire Usal, Laurent Tesson, Mathieu Rouel, Flore Couzy, Francine Padonou, Cloé Zamit, Jeremy Santamaria, Pierre Maminirina, Olivier Baron, Jerome Jullien, Jérémie Poschmann, Magali Irla, Ignacio Anegon, Matthieu Giraud

## Abstract

Promiscuous expression of tissue-specific antigens (TSAs) by medullary thymic epithelial cells (mTECs) underpins central T-cell tolerance, relying on NF-κB-driven mTEC maturation and induction of the autoimmune regulator AIRE. However, a substantial fraction of TSAs is AIRE-independent, implying additional regulators that remain only partially identified. Using cross-species single-cell chromatin accessibility profiling of human and mouse TECs, we find that HIVEP motifs are among the most accessible regulatory elements across the mTEC lineage. Among HIVEP paralogs, *HIVEP3* is robustly expressed in mature mTECs across human, mouse and rat; given that rat immunity more closely mirrors human physiology than the mouse, we generated a CRISPR *Hivep3*-knockout rat line. *Hivep3* deletion impairs mTEC maturation, reduces *Aire* expression and reshapes the TSA repertoire that underlies negative selection and Treg induction, as Hivep3 constrains canonical NF-κB1 and sustains non-canonical NF-κB2, the principal mediator of mTEC maturation. Beyond mTECs, *Hivep3* deficiency downregulates Foxn1 target genes in cortical TECs and impairs positive selection. In the periphery, *Hivep3*-KO rats show reduced splenic CD4⁺ T-cells, a shift from naïve towards effector-and central-memory phenotypes, increased regulatory T-cells, and a sustained Th1-biased serum profile (elevated IFN-γ, reduced IL-17A) with concurrent systemic metabolic perturbations across all ages. With age, this culminates in chronic inflammation, with multi-organ CD3⁺ T-cell infiltration accompanied by inflammatory lesions. Together, these findings establish HIVEP3 as a previously unrecognised regulator of thymic epithelial function and central tolerance, whose deficiency leads to systemic T-cell-mediated chronic inflammation.

## INTRODUCTION

The thymus is a primary lymphoid organ essential for generating a functionally diverse and self-tolerant T-cell repertoire, enabling protection against infection and tumours while avoiding autoimmunity^1,2^. To achieve this, immature thymocytes undergo successive rounds of selection within a stromal microenvironment governed by the transcription factor Foxn1, which maintains thymic epithelial cell (TEC) identity and competence^3^. First, cortical TECs (cTECs), through their unique molecular machinery, drive thymocyte lineage commitment and positive selection in the cortex, thereby shaping the primary T-cell repertoire and ensuring recognition of self-MHC molecules with sufficient affinity^3,4^. Subsequently, selected thymocytes migrate to the medulla, where medullary thymic epithelial cells (mTECs), along with other antigen-presenting cells (APCs) such as dendritic cells and B cells, mediate the negative selection of autoreactive cells^5^. mTECs are uniquely poised to express thousands of genes that are typically restricted to peripheral tissues, known as tissue-specific antigens (TSAs), enabling a broad spectrum of self-antigens to encounter developing thymocytes^6,7^. This results in the deletion of autoreactive clones or their conversion into regulatory T-cells (Tregs), thereby preventing autoimmunity.

In the thymus, interactions with developing thymocytes promote mTEC maturation, with the expression of high levels of MHC-II molecules and a broad repertoire of TSAs. The development and maintenance of mTECs are tightly regulated by tumour necrosis factor (TNF) superfamily signalling initiated by RANK and CD40, which engage both the canonical and the non-canonical NF-κB pathways^8^. The non-canonical pathway is the principal driver of mTEC maturation and AIRE induction: NF-κB-inducing kinase (NIK) mediates proteolytic processing of the NF-κB2 precursor p100 into its active form p52, allowing nuclear translocation of the RelB/p52 complex^9^. In parallel, the canonical pathway leads to nuclear translocation of the RelA/p50 complex (NF-κB1) and is classically associated with the control of inflammatory gene programs; how their respective outputs are balanced in mTECs remains incompletely understood.

The transcriptional regulator AIRE is a key driver of TSA expression in the thymus. In contrast to traditional transcription factors (TFs), which bind specific DNA sequences, AIRE preferentially targets repressed genes, including TSAs, by reading unmethylated histone H3K4 (H3K4me0) through its PHD1 domain, a mark of transcriptionally silent chromatin^10^. In humans, loss-of-function mutations in *AIRE* lead to a rare monogenic autoimmune disease, autoimmune polyendocrinopathy-candidiasis-ectodermal dystrophy (APECED), characterised by multi-organ autoimmunity, including type 1 diabetes and adrenal insufficiency, highlighting the direct link between mTEC-intrinsic defects and systemic autoimmunity^11,12^. Notably, a prominent Th1/IFN-γ component is a recurring feature of both organ-specific autoimmune diseases and chronic inflammatory conditions, yet how TEC-intrinsic defects shape peripheral Th-cell polarisation remains unclear^13^.

Single-cell technologies have significantly advanced our understanding of mTEC diversity and TSA expression, revealing that the mTEC compartment extends beyond AIRE-expressing cells to encompass a heterogeneous population of mimetic mTECs^14^. These cells express TSAs in a biologically coherent manner through lineage-specific TFs (e.g., Neurod1 for neuroendocrine, Foxa1/2 for lung, Sox8 for microfold and Myog for muscle)^14^. While some mimetic populations are modulated by AIRE, such as FoxA^+^ and Sox^+^ cells, others develop in its absence, indicating that the regulatory landscape governing TSA expression is more complex than previously appreciated^14^. Despite these advances, the transcriptional regulators that drive mTEC maturation and TSA expression within the *AIRE*-expressing compartment itself remain incompletely resolved.

To uncover transcriptional regulators operating in mature mTECs beyond AIRE, we leveraged cross-species single-cell chromatin accessibility profiling of human and mouse thymic epithelial cells. This unbiased approach identified HIVEP recognition motifs as being highly enriched in *AIRE*-accessible mTECs. HIVEP proteins are a family of three large zinc-finger TFs that bind κB-like sequences, but whose role in TEC biology has not been explored^15^. Among the three paralogs, *HIVEP3* was consistently expressed in human, mouse and rat mTEC^AIRE^. To dissect its function *in vivo*, we generated the first *Hivep3*-knockout rat model. Herein, we demonstrate that in rats, Hivep3 drives mTEC maturation through constraining canonical NF-κB1 activity and promoting non-canonical NF-κB2 signalling, sustains both AIRE-dependent and AIRE-independent TSA expression, and shapes the cTEC transcriptional network controlling thymocyte positive selection. The resulting compromised central tolerance manifests in the periphery as Th1-driven chronic systemic inflammation, thereby establishing HIVEP3 as a central regulator of thymic function and immune homeostasis.

## RESULTS

### The regulatory framework of human TECs uncovers TFs driving maturation and mimetic cell fate

In humans, to investigate genome-wide regulatory activity underlying mTEC heterogeneity, we performed single-cell ATAC sequencing (scATAC-seq) on four paediatric thymic samples removed during cardiothoracic surgery. The four donors included two females (7 and 13 days) and two males (7 and 12 days). mTECs were enriched through sequential magnetic separation, first depleting CD45^+^ cells, then selecting epithelial cells with EpCAM. This pre-enriched population underwent further EPCAM⁺CD205⁻HLA-DR⁺ isolation via fluorescence-activated cell sorting (FACS) to select MHC-II^+^ mTECs and deplete the majority of cTECs. After standard pre-processing and batch-effect correction, dimensionality reduction followed by unsupervised genome-wide chromatin accessibility clustering allowed us to identify nine TEC clusters (Figure 1A *Top*; gating strategy in Figure S1A). These clusters reflected biological heterogeneity rather than batch or technical variation, as supported by the overlapping distribution of donors across clusters (Figure 1A *Bottom*) and by the consistent quality metrics across all four donors, including strong transcription start site (TSS) enrichment, high fraction of reads in peaks, and elevated numbers of unique nuclear fragments per cell (Figure S1B). Cells with open chromatin at *AIRE* and MHC-II gene loci (*HLA-DRA, HLA-DQA1, HLA-DRB1*) were annotated as mTEC^AIRE^ (Figure 1B). Three mTEC clusters were defined as mimetic mTEC^Keratinocyte^ (*IVL*), mTEC^Neuroendocrine^ (*NEUROD2*) and mTEC^Muscle^ (*MYOD1*) based on chromatin accessibility at lineage-defining factor loci. Interestingly, the mTEC^Keratinocyte^ cluster showed substantial chromatin accessibility at the *AIRE* locus and was positioned next to mTEC^AIRE^, consistent with a direct divergence from the *AIRE*-expressing state. The identity of the mTEC^AIRE^ and these three mimetic clusters was confirmed by projecting the re-analysed human scRNA-seq data from *Huisman et al.*, 2025^16^ (Figure 1C). This projection also enabled the identification of a Transitional mTEC cluster adjacent to the mTEC^Neuroendocrine^ and mTEC^Muscle^ clusters, for which no clear lineage-defining factor loci could be identified (Figure 1B), as well as a mimetic mTEC^Ionocyte^ cluster (Figure 1C).

**Figure 1:**
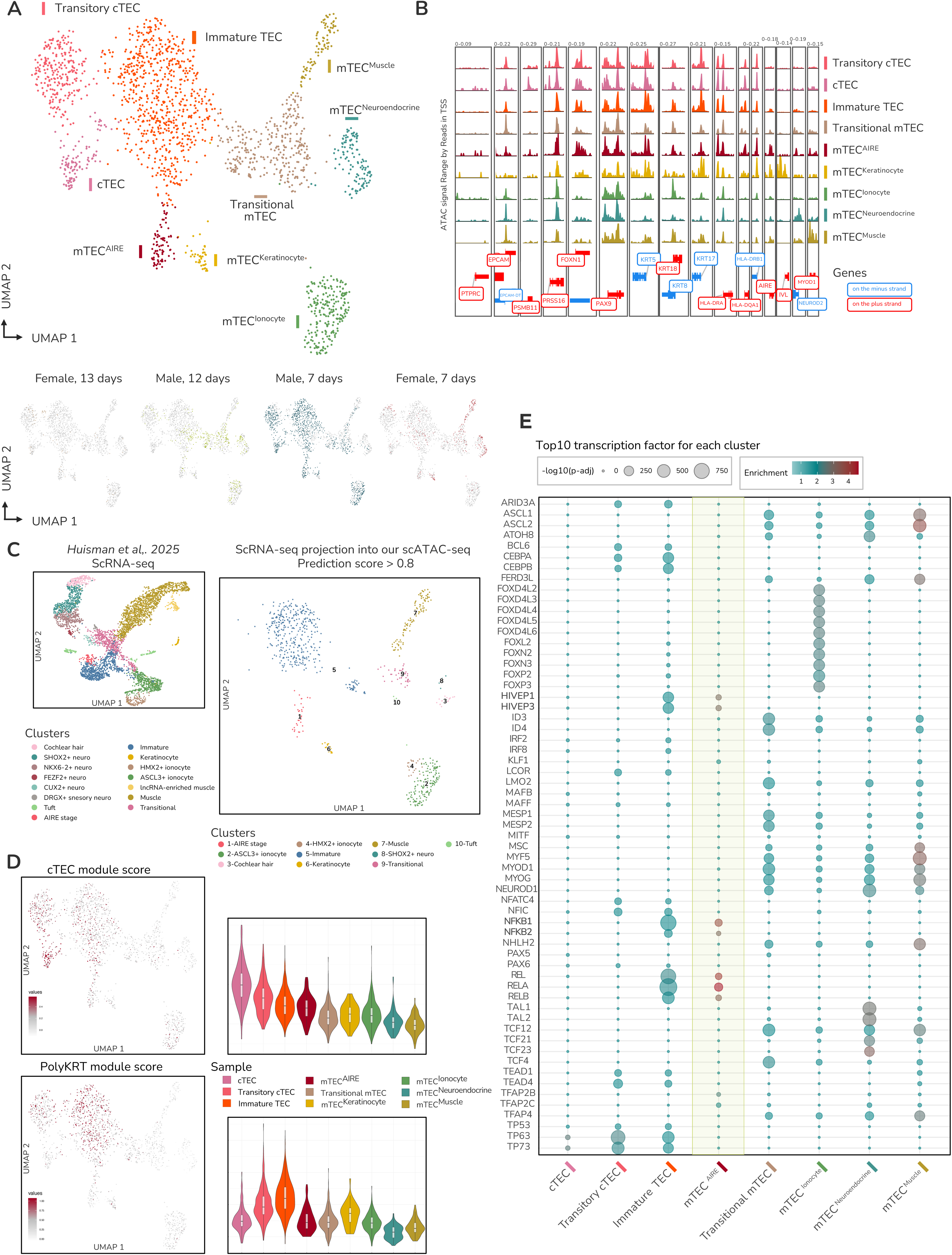
The regulatory architecture of human TEC reveals the transcriptional drivers of TEC maturation and subset identity. **(A)** Uniform manifold approximation and projection (UMAP) of human scATAC-seq. A total of 9 clusters were resolved and colour-coded. Each dot is a single nucleus. (top) UMAP representation of 4 thymic donors resolved by colour (bottom). **(B)** Chromatin-accessibility tracks at indicated loci. Genes localised on the minus strand are in blue while genes on the plus strand are in red. **(C)** Projection of scRNA-seq data from *Huisman et al., 2025*. On the left, UMAP of reanalysed scRNA-seq. Each dot is a single cell. On the right, UMAP of the scATAC-seq with only the nuclei presenting a prediction score higher than 0.8. The predicted clusters are annotated and colour-coded. **(D)** Module score analysis of cTEC (top) (*SCX, CFC1, PSMB11, TP53AIP1, SLC46A2, LY75, KCNIP3, FOXN1, CD274, TBATA, PRSS16, CTSV, CCL25*) and polyKRT populations (bottom) (*KRT13, KRT14, KRT15, KRT17, KRT19, CCL19, CEBPD, CLU, FN1, IFITM3, VCAM1, TAGLN, BCAM, LIFR,* and *CH25H*). Module scores are represented on the UMAP (left) and in the violin plot (right). **(E)** Comparative analysis of the top 10 enriched motifs across each cluster using the CIS-BP database for Homo sapiens. The transcription factors are ordered alphabetically. The size of the dot represents-log10(adjusted p-value), while the colour indicates the enrichment score.

We then showed that the three remaining clusters shared increased chromatin accessibility at *FOXN1*, *PAX9*, and at several keratin-encoding gene loci (*KRT5, KRT8, KRT17, KRT18*) (Figure 1B). Two of them were further characterised by accessibility at the *PRSS16* locus, indicating cTEC identity, whereas the third corresponds to Immature TEC as shown by the human scRNA-seq projection (Figure 1C). Therefore, we note an incomplete cTEC depletion in the dataset, despite the CD205⁻ gating strategy. In addition, the two cTEC clusters showed differential accessibility at *PSMB11,* suggesting different maturation states (Figure 1B). A module score analysis of a mature cTEC signature (*SCX, CFC1, PSMB11, TP53AIP1, SLC46A2, LY75, KCNIP3, FOXN1, CD274, TBATA, PRSS16, CTSV, CCL25*) showed high levels in the *PSMB11*-accessible cluster and lower levels in the non-accessible one (Figure 1D, *Top*), allowing their characterisation as mature and transitory cTEC clusters, respectively. Regarding the Immature TEC cluster, we took advantage of the recent dataset by *Ragazzini et al., 2023*, in which they identified a human immature TEC subset named “*polyKRT*”due to its co-expression of multiple keratin genes^17^. This population functions as an adult TEC stem/progenitor population capable of generating both mTECs and cTECs. To further evaluate whether our Immature TEC cluster aligns with the polyKRT one, we examined the accessibility of the polyKRT-associated gene signature defined by these genes (*KRT13, KRT14, KRT15, KRT17, KRT19, CCL19, CEBPD, CLU, FN1, IFITM3, VCAM1, TAGLN, BCAM, LIFR,* and *CH25H*), and found that it is higher in the Immature TEC, indicating that this cluster may represent human bipotent immature TECs (Figure 1D, *Bottom*).

Then, to further delineate TFs underlying the mTEC clusters, we performed cluster-level TF motif enrichment using the *Homo sapiens* CIS-BP database on accessible regions defined by stringent peak selection criteria (|log₂FC| ≥ 0.5, FDR ≤ 0.1) (results shown in Table 1). This analysis identified NF-κB family motifs (NF-κB1, NF-κB2, REL, RELB) as the most prominently enriched motifs in mTEC^AIRE^, consistent with prior studies in mice demonstrating the central role of NF-κB, notably NF-κB2, in driving *Aire* expression and MHC-II^high^ mTEC development^18,19^. Beyond NF-κB, HIVEP family motifs (HIVEP1, HIVEP3) ranked second, identifying regulatory candidates not previously implicated in mTEC biology. HIVEP proteins have been reported to bind κB-like sequences, often overlapping with NF-κB recognition motifs, and to directly compete with NF-κB for DNA occupancy^20,21^ (Figure S1C). Beyond their TF features, HIVEP proteins can also function as adapter proteins^15,22,23^. Interestingly, we also found that HIVEP motifs were strongly enriched in the Immature TEC cluster, suggesting that HIVEP sustained a regulatory program that is already in place in bipotent progenitors.

Regarding mimetic TECs, the enriched motif analysis revealed both shared and unique TF signatures. mTEC^Ionocyte^ was uniquely enriched for FOX motifs and did not overlap with other clusters, suggesting a distinct regulatory program (Figure 1E). In contrast, the mTEC^Neuroendocrine^ and mTEC^Muscle^ clusters shared enrichment for multiple basic helix-loop-helix (bHLH) family factors, including MYOD, MYOG, and NEUROD1, likely reflecting a permissive chromatin state enabling cross-lineage TF binding and supporting broad TSA representation.

Collectively, we establish a chromatin landscape atlas of human TECs that uncovers the regulatory programs underlying TEC heterogeneity and maturation. Notably, our analysis reveals the early engagement of an NF-κB and HIVEP regulatory axis in mTEC maturation, alongside distinct and overlapping TF signatures in differentiated mimetic cells.

### Chromatin accessibility profiling of mouse TECs uncovers signatures underlying distinct mTEC lineages

Having established the regulatory TF network of human TECs and identified NF-κB and HIVEP TF families as prominently enriched in Immature TEC and mTEC^AIRE^, we next sought to determine whether these findings are evolutionarily conserved by performing parallel scATAC-seq analyses in mice. To this end, we performed scATAC-seq on CD45⁻EpCAM⁺ TECs isolated by sequential magnetic separation from pooled thymi of 4-week-old mice. After quality filtering based on TSS enrichment and fragment count, we predominantly retained high-quality mTEC nuclei with minimal thymocyte carryover. As for humans, the mouse dataset exhibited strong nucleosomal patterning and high TSS signal enrichment, enabling the identification of 13 distinct clusters based on chromatin accessibility profiles (Figures 2A, S2A). In contrast to the carryover thymocytes, all other clusters exhibited high accessibility at loci encoding the epithelial marker *Epcam* and major MHC-II genes, including *H2-Ab1, H2-Aa, and H2-Eb1* (Figure 2B). Notably, three clusters displayed open chromatin at the *Aire* locus, including at *Aire* known cis-regulatory elements^24^. We grouped these clusters as “*Aire-*accessible mTECs” and refined their characterisation by projecting a high-coverage mouse scRNA-seq dataset, which we generated, capturing the full spectrum of mTEC maturation states (Figures 2B, S2B). This approach enabled the assignment of the three clusters as i) transit-amplifying TECs (TAC-TECs), precursors of both Ccl21^+^ and *Aire*-expressing mTECs ^25^, ii) mTEC^Aire^ ^Early^ and iii) mTEC^Aire^ ^Late^.

**Figure 2:**
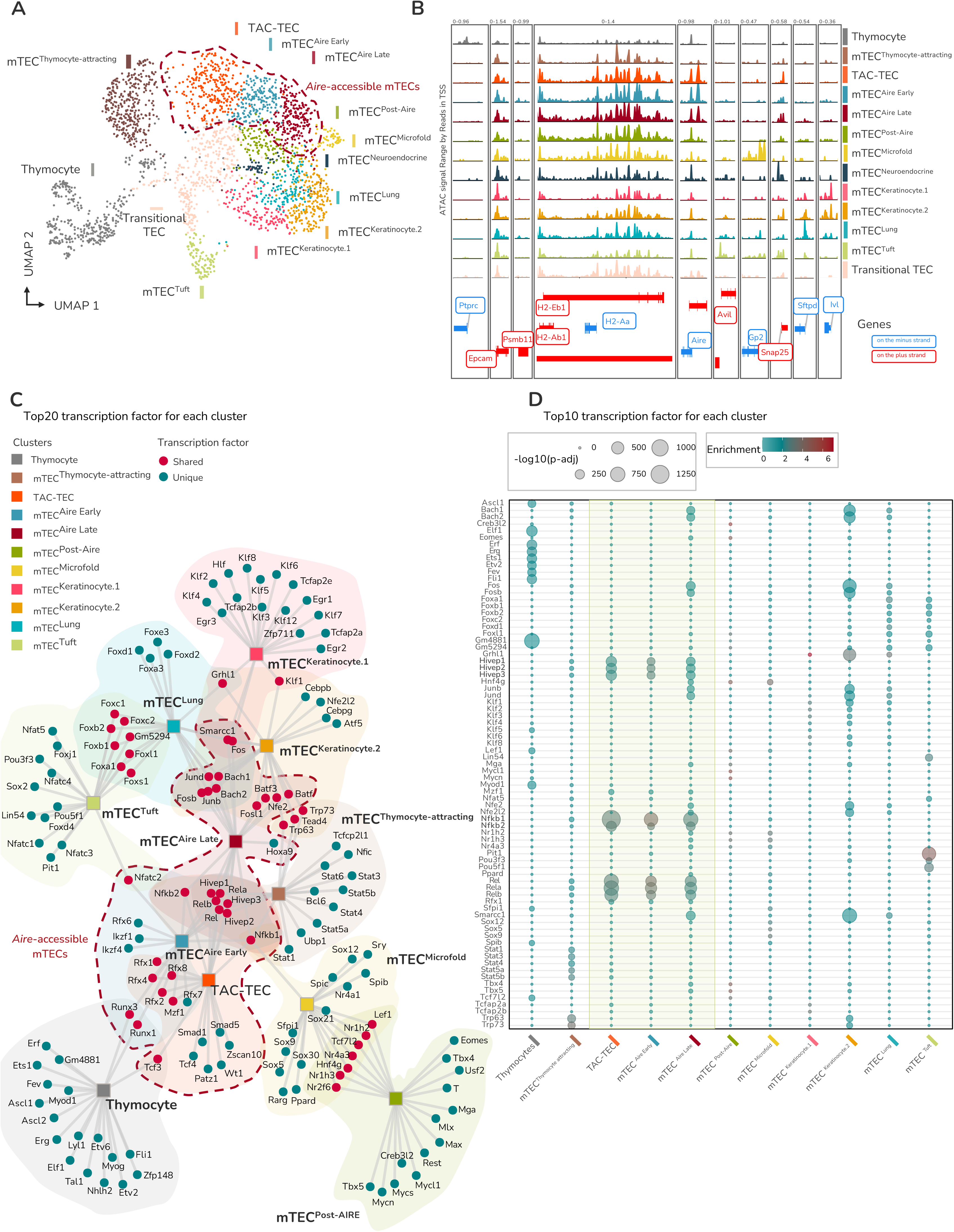
scATAC-seq reveals chromatin signatures underlying distinct mTEC lineages in the mouse. **(A)** Uniform manifold approximation and projection (UMAP) of scATAC-seq of 2,573 nuclei. A total of 13 clusters were resolved and colour-coded. Each dot is a single nucleus. **(B)** Chromatin-accessibility tracks at indicated loci. Genes localised on the minus strand are in blue while genes on the plus strand are in red. **(C)** String plot of the top 20 enriched transcription factor motifs in each cluster. The clusters are indicated in squares, coloured by the indicated cluster. The transcription factor is represented by a circle; circles are blue when unique and pink when shared with other clusters. The “*Aire*-accessible mTEC” is framed in red. **(D)** Comparative analysis of the top 10 enriched motifs across each cluster using the CIS-BP database for Mus musculus. The transcription factors are ordered alphabetically. The size of the dot represents-log10(adjusted p-value), while the colour indicates the enrichment score.

In contrast, *Aire* regulatory elements were not accessible in the remaining clusters. Most of them exhibited accessible regions at genes known to drive various mimetic programs, as described in previous studies ^26,27,14^. Hence, we could identify the tuft-like mTEC cluster marked by *Avil* accessibility, alongside additional mimetic subtypes corresponding to microfold (*Gp2*), neuroendocrine (*Snap25*), lung-associated (*Sftpd*), and keratinocyte-like (*Ivl*) mTECs^28,29^. Their mimetic identity was confirmed by the projection of our mouse scRNA-seq dataset (Figures 2B, S2B). We further observed epigenetic heterogeneity within the keratinocyte-like lineage, resolving it into two distinct sub-clusters. A cluster adjacent to mTEC^Aire^ ^Late^ and to neighbouring mimetic subsets, lacking clear lineage-defining open loci, was generically labelled mTEC^Post-Aire^. A separate, centrally positioned cluster, similarly devoid of defining open loci, was designated as Transitional TEC. Finally, the last cluster, further apart, was found to correspond to the Ccl21^+^ mTEC population (Figure S2B), and was labelled mTEC^Thymocyte-attracting^ as *Ccl21a*, *Ccl21b* and *Ccl21c* loci were absent from the used genome annotations.

### Shared and lineage-specific regulatory signatures sustain mTEC maturation and mimetic differentiation

As in the human dataset, we performed cluster-level TF motif enrichment using the cis-BP database for *Mus musculus*, on accessible regions defined by stringent peak selection criteria (|log₂FC| ≥ 0.5, FDR ≤ 0.1) (results shown in Table 2). First, among the three “*Aire*-accessible mTEC” clusters, TAC-TEC, mTEC^Aire^ ^Early^ and mTEC^Aire^ ^Late^, we observed both shared and cluster-specific TF-motif enrichment, indicating that distinct TFs operate at successive stages of mTEC maturation (Figures 2C-D). Consistent with prior studies and with our human scATAC-seq data, the most prominently enriched motifs across these clusters belonged to the NF-κB family (NF-κB1, NF-κB2, Rel, Relb), closely followed by HIVEP family members (Hivep1, Hivep2, Hivep3), which recognise closely related but slightly divergent κB-like sequences (Figure S2C). Of note, in mice, the binding motifs of all HIVEP family members were inferred from the human HIVEP2 motif. Collectively, the co-enrichment of NF-κB and HIVEP motifs, particularly from the TAC-TEC stage onward, indicates that the HIVEP regulatory program is engaged early in mTEC differentiation. This suggests that HIVEP factors may modulate NF-κB-dependent gene regulatory programs, either by competing for shared κB binding sites or by fine-tuning downstream transcriptional output, thereby influencing mTEC maturation and TSA expression.

Within mimetic mTECs, each cell type exhibited accessible chromatin regions enriched for lineage-specific TF motifs, including Grhl1 for keratinocyte-like mTEC, Spib along with SOX family motifs for mTEC^Microfold^, POU family motifs for mTEC^Tuft^ and FOXA family motifs for mTEC^Lung^. While some of these motifs were exclusive to their associated cluster, and others ranked among the top 20 significantly enriched factors in additional mimetic clusters (Figure 2C), they all reached the highest statistical significance in their associated cluster (Figure 2D), supporting these lineage-specific assignments. Interestingly, cluster-specific motif comparisons revealed associations between motif enrichment patterns and mTEC maturation states. Notably, the mTEC^Aire^ ^Late^ cluster exhibited a hybrid motif signature, retaining early-stage motifs, such as those from the NF-κB family, while acquiring those linked to post-Aire or terminal differentiation, including Hnf4g (ranked 23^rd^) and Smarcc1 (ranked 6^th^), a core component of the SWI/SNF chromatin-remodelling complex (Figures 2D). Together, these observations suggest that the transition from early *Aire*-accessible mTECs to specialised mimetic mTECs may occur through a gradual and coordinated reorganisation of TF networks, rather than an abrupt regulatory shift, reflecting the layered complexity of mTEC maturation and central tolerance.

### HIVEP motif enrichment during mTEC maturation is conserved across human and mouse *AIRE*-accessible mTECs

We next investigated cross-species conservation of TF motif enrichment in mTECs by comparing our mouse and human scATAC-seq datasets, focusing on *AIRE*-accessible mTEC clusters (Figure 3A-B). Examination of enriched motifs shared in human and mouse revealed 36 shared TFs, with a pronounced co-enrichment of NF-κB and HIVEP motifs across genome-wide accessible regions (Figure 3B), supporting a model in which HIVEP factors contribute to mTEC maturation and TSA regulation.

**Figure 3:**
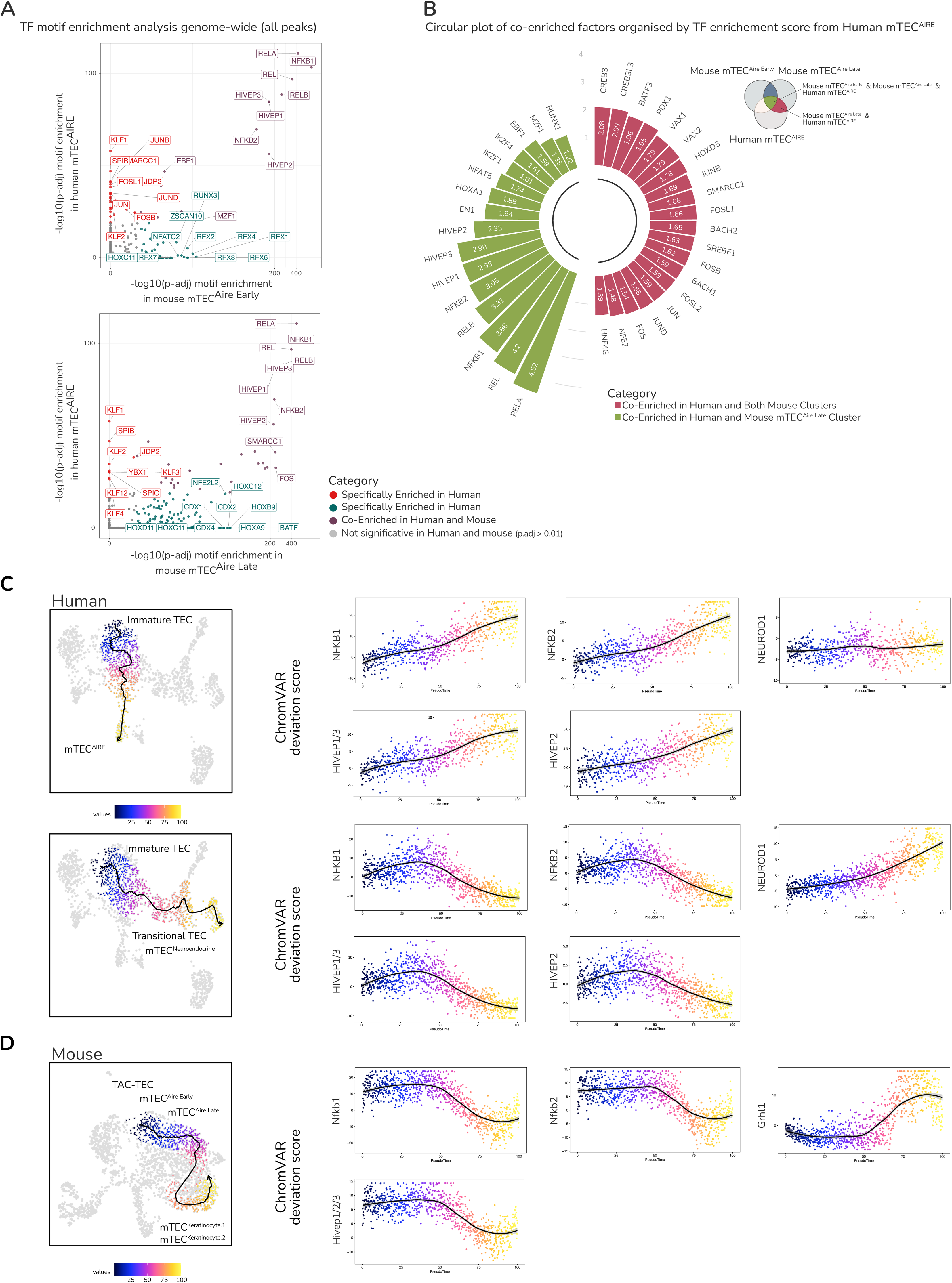
The HIVEP transcription factor family is the most evolutionarily conserved transcription factor family, along with NF-κB, in *Aire*-expressing mTECs. **(A)** Scatter plot showing enrichment scores (-log10(adjusted p-value) in mouse mTEC^Aire^ ^Early^ and mTEC^Aire^ ^Late^ compared to human mTEC^AIRE^ cluster. This analysis was performed using all peaks. Each dot represents a transcription factor and is coloured based on the type of enrichment. **(B)** Circular bar plot of the 36 TFs shared between *Aire*-accessible human and mouse clusters. The bar plot is organised based on human mTEC^AIRE^ TF enrichment scores, with the scores annotated within the bars. Colour represents the type of co-enrichment: green if enriched in human mTEC^AIRE^ and both mouse *Aire*-accessible mTEC clusters; red if enriched in human mTEC ^AIRE^ and only mouse mTEC ^AIRE^ ^late^. **(C)** ChromVAR deviation score trajectory in human scATAC-seq data spanning immature and mTEC^AIRE^ (top) or from immature and transitional mTEC to mTEC^Neuroendocrine^ for NFKB1/2, HIVEP1/2/3 and NEUROD1 factors. **(D)** ChromVAR deviation score trajectory in mouse scATAC-seq data spanning TAC-TEC, mTEC^AireEarly^, mTEC^Aire^ ^Late^, mTEC^Keratinocyte.1^ and mTEC^Keratinocyte.2^ with individual feature plots for Nfkb1/2, Hivep1/2/3 and Grhl1 genes. Dot plots show the ChromVAR deviation score for the corresponding transcription factors.

To gain insight into the regulatory dynamics of HIVEP factors during mTEC development, we analysed NF-κB1 and NF-κB2 alongside the HIVEP family across pseudotime-ordered mTEC clusters using chromVAR deviation scores, which reflect TF binding motif accessibility variability, and gene activity scores, which serve as a proxy for chromatin accessibility around gene bodies (Figure S3A-B). In the human dataset, on the basis of UMAP positioning, we examined two trajectories from Immature TEC: one to the adjacent mTEC^AIRE^, and another to the mimetic mTEC^Neuroendocrine^ through Transitional mTEC. We found that NF-κB1, NF-κB2 and HIVEP family binding motifs (shared HIVEP1/3 motif in human) were selectively enriched along the Immature TEC to mTEC^AIRE^ trajectory, whereas they were progressively lost along the other trajectory toward mTEC^Neuroendocrine^. In both trajectories, gene activity scores at the *NFKB1, NFKB2*, *HIVEP1*, *HIVEP2* and *HIVEP3* loci remained relatively uniform, consistent with these loci remaining accessible to transcriptional regulation throughout mTEC maturation (Figure 3C, S3A). By contrast, NEUROD1, a lineage-defining TF for neuroendocrine-like mTECs, showed both strong motif enrichment and elevated gene activity scores confined to its corresponding mimetic cluster, mTEC^Neuroendocrine^, consistent with a direct role in establishing subset-specific identity. When we applied the same analytical framework to the mouse dataset, spanning a trajectory from TAC-TEC through mTEC^Aire^ ^Early^ and mTEC^Aire^ ^Late^ to mTEC^Keratinocyte.1^ and mTEC^Keratinocyte.2^, with Grhl1 serving as the lineage-defining TF for keratinocyte-like mTECs, we observed a remarkably comparable pattern. NF-κB and Hivep family (shared HIVEP family motif in mouse) motif enrichment was high in TAC-TEC and mTEC^Aire^ clusters and dropped in mimetic mTEC clusters, while gene activity scores at all three *Hivep* loci remained stable along the trajectory (Figure 3D, S3B). Together, in humans and mice, *NFKB* and *HIVEP* family loci remained accessible to transcriptional regulation throughout mTEC maturation, while HIVEP and NF-κB binding activity was enriched at early-to-intermediate stages and declined as cells acquired mimetic fates. This conserved temporal pattern positions the HIVEP–NF-κB axis as an early driver of mTEC maturation, distinct from lineage-defining TFs that act within mimetic subsets.

### Cross-species expression profiling identifies *HIVEP3* as a conserved *HIVEP* paralog in *AIRE*-expressing mTECs

Because motif analysis cannot discriminate between HIVEP paralogs, we examined the expression pattern of each HIVEP family member (*HIVEP1*, *HIVEP2* and *HIVEP3*) in scRNA-seq datasets across TEC clusters. In humans, in the re-analysed scRNA-seq dataset from *Huisman et al., 2025*, all three *HIVEP* paralogs displayed robust expression in AIRE-positive mTECs, and broad distribution across other mTEC subsets, including mimetic cells, which precluded clear discrimination between paralogs (Figure 4A). In our mouse scRNA-seq dataset, *Hivep1* and *Hivep3* were both expressed in Aire-positive mTECs, unlike *Hivep2,* which was detected only at very low levels within this compartment (Figure 4B). Given the lack of discriminatory expression between *Hivep1* and *Hivep3* transcripts, we extended our analysis to the rat using the scRNA-seq dataset from *Stoljar et al., 2025*. Here, we found that *Hivep3* was the most highly expressed HIVEP family member in Aire-positive mTECs, defined as mTEC II, also referred to as mTEC^High^. Indeed, as in mice, *Hivep2* was very low, while *Hivep1* appeared lower than in mice in this dataset, yielding a higher *Hivep3*/*Hivep1* expression ratio (Figure 4C). We therefore selected the rat, primarily because its immunity more closely mirrors human physiology than the mouse, with the higher Hivep3/Hivep1 ratio further suggesting a reduced risk of paralog compensation. Of note, *Hivep3* expression was also observed in rat cTECs, the latter being well captured in the used scRNA-seq dataset, in contrast to the human and mouse ones.

**Figure 4:**
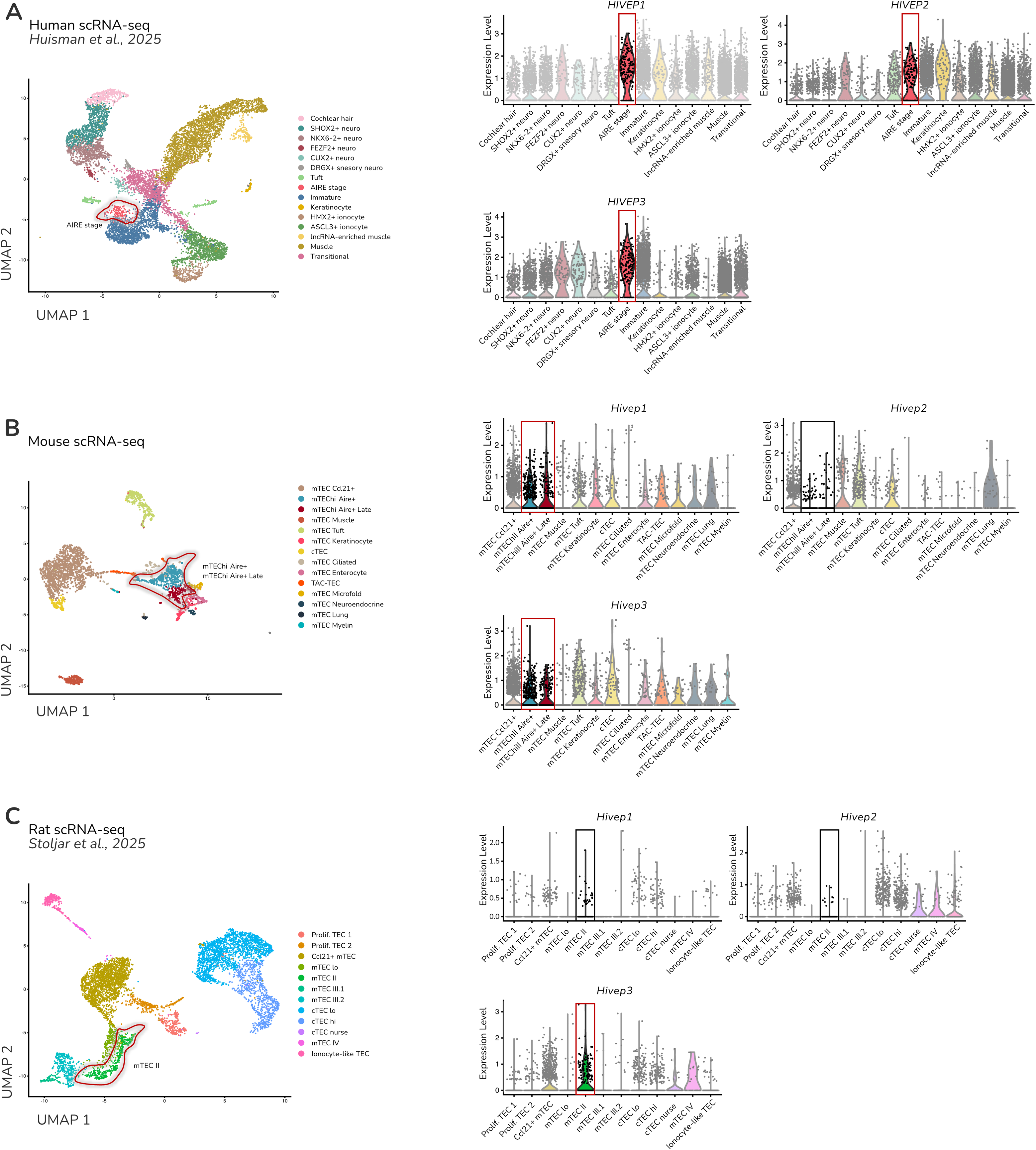
HIVEP3 is consistently expressed in *AIRE*-expressing mTECs across human, mouse and rat. Uniform manifold approximation and projection (UMAP) of mTEC clusters (left) and violin plot showing the expression of *HIVEP1*, *HIVEP2* and *HIVEP3* from the re-analysed scRNA-seq dataset from **(A)** human (*Huisman et al., 2025*), **(B)** mouse (this study), and **(C)** rat (*Stoljar et al. 2025*). Each dot represents a cell. *Aire*-accessible clusters are framed in red.

### *Hivep3* deficiency disrupts thymic architecture and reduces Aire expression

Therefore, to investigate the role of HIVEP3, we generated a CRISPR/Cas9-mediated knockout (KO) in Sprague-Dawley rats by introducing frameshift mutations in the first exon of the *Hivep3* gene using the genome-editing via oviductal nucleic acids delivery (GONAD) method, an *in vivo* electroporation-based approach that delivers CRISPR components directly into the oviduct (Figure 5A). KO efficiency and loss of *Hivep3* transcript were subsequently confirmed by RT-PCR in the thymus and brain, the latter being a tissue where *Hivep3* is known to be highly expressed (Figures 5A-S4A). Consistent with its established role in bone biology, *Hivep3* deletion in rats enhanced osteogenesis and reduced bone marrow, mirroring observations in mouse models^30^ (Figure S4B).

**Figure 5:**
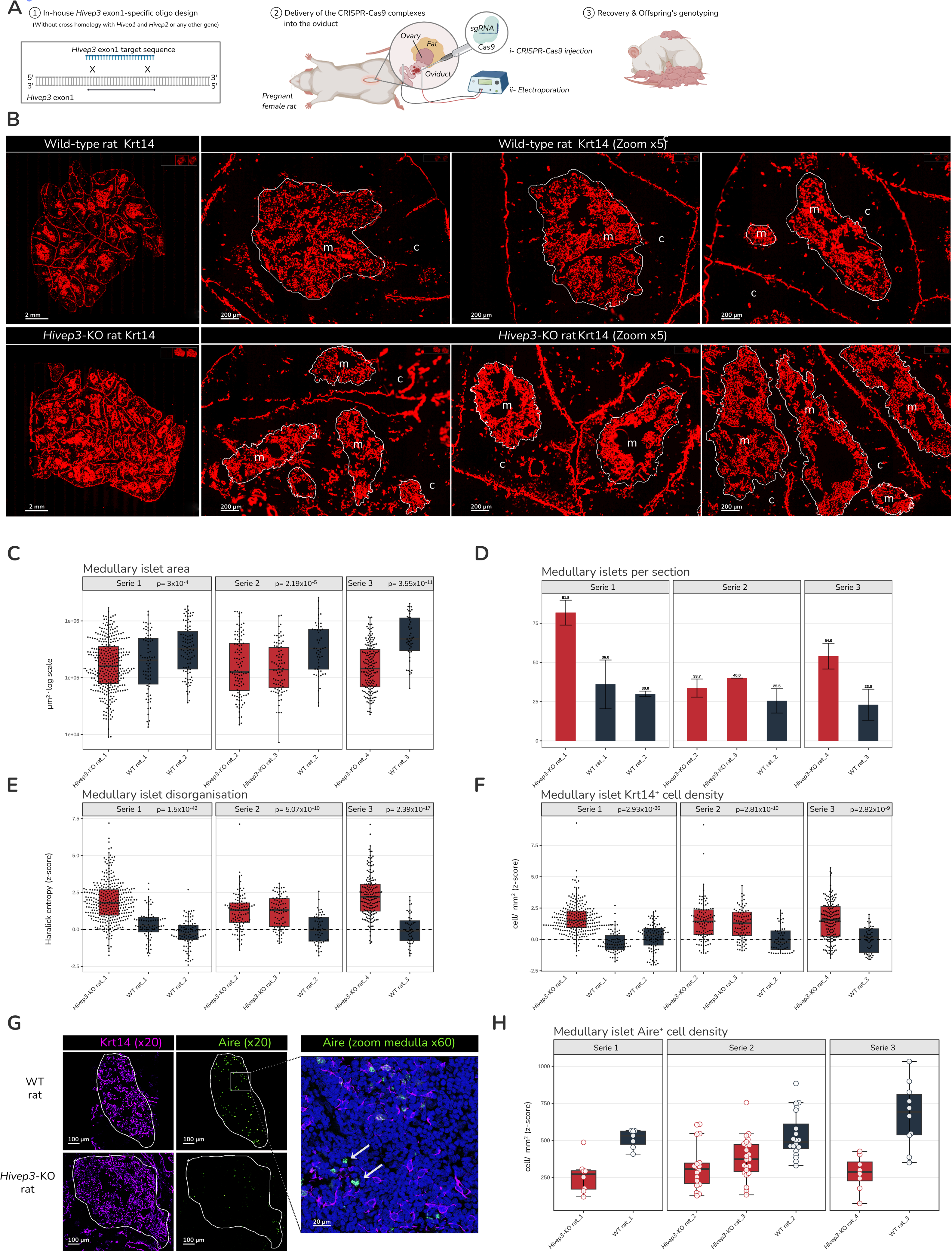
Hivep3 deficiency disrupts thymic architecture and perturbs Aire expression. **(A)** Schematic representation of the generation of the *Hivep3*-deficient rat model. **(B)** Immunofluorescence microscopy of WT and *Hivep3*-KO thymic sections stained for Krt14. Scale bar, 2 mm (left) and a zoom of the medullary section of the thymus (right). m:medulla, c:cortex. Bar plot of **(C)** log₁₀-transformed medullary surface area (μm²), **(D)** average medullary islet counts, (**E)** Haralick entropy (Z-score) and **(F)** density of Krt14^+^ cells per mm² (Z-score). **(G)** Immunofluorescence microscopy of WT and *Hivep3*-KO thymic sections stained for Krt14 (violet) and Aire (green). Scale bar, 100 μm (left) and a zoom at medulla with an arrow pointing at AIRE staining. Scale bar, 20 μm (right). **(H)** Bar plot of the number of Aire-positive cells per mm². *Hivep3*-KO condition is represented in red, while the WT is in blue.

We then assessed the impact of *Hivep3* deficiency on thymic architecture via immunofluorescence (IF) staining for Krt14, a known marker of mTECs, on thymic sections from *Hivep3*-KO and WT rats. In *Hivep3*-KO thymi, we observed more fragmented and less cohesive medullary islets compared to WT controls (Figure 5B). Quantitative image analysis corroborated this observation, revealing a significantly reduced medullary islet area (Figure 5C), a higher number of medullary islets per thymic section (Figure 5D), and increased structural disorganisation of medullary islets, as measured by Haralick entropy (Figure 5E). In addition, we observed a significant increase in Krt14⁺ cell density within medullary islets of *Hivep3*-KO thymi (Figure 5F). By re-analysing the *Park et al., 2020* human thymic atlas, we previously showed that KRT14 preferentially marks early-stage mTECs^31,32^. Consistent with this, using the *Stoljar et al., 2025* scRNA-seq dataset from WT rat, we found that *Krt14* expression is largely absent from mTEC^High^ and most prominent in Ccl21⁺ and immature TEC populations (Figure S4C). Consequently, the increased density of KRT14⁺ cells in *Hivep3*-KO medullary islets may reflect a failure or arrest in mTEC maturation, which could underlie the disrupted thymic architecture observed in these rats.

Given that the transcriptional regulator AIRE is a key marker of mTEC maturation and a major inducer of TSA expression, we next examined AIRE expression by IF within medullary islets of *Hivep3*-KO and WT rats (Figure 5G). Consistent with an mTEC maturation defect, Aire⁺ cell density was significantly reduced in *Hivep3*-deficient thymi compared to WT (Figure 5G-H). Taken together, these findings indicate that Hivep3 is required for proper progression along the mTEC maturation program, and its loss may, directly or indirectly, impact TSA expression.

### Hivep3 constrains canonical NF-κB1 and sustains non-canonical NF-κB2 signalling in mTEC^High^

To gain insight into the molecular mechanisms underlying the mTEC maturation defect observed in *Hivep3*-KO rats, we performed scRNA-seq on pooled thymi from *Hivep3*-KO and WT rats, focusing on the TEC compartment (Figure 6A, S5A). Consistent with our IF observations, differential expression (DE) analysis confirmed significant downregulation of *Aire* in *Hivep3*-KO mTEC^High^ (Figure 6B; Table 3). Unexpectedly, in contrast to the *Stoljar et al. 2025* dataset, where *Hivep1* expression in rat mTEC^High^ was minimal (Figure 4C), our WT dataset revealed comparable expression levels of *Hivep1* and *Hivep3* (Figure S5B). This discrepancy may reflect genetic variation between outbred Sprague-Dawley colonies. Similar to our observations in the mouse, this raises the possibility of functional redundancy between Hivep3 and Hivep1. Nevertheless, *Hivep1* expression remained unchanged between genotypes (Figure S5B), indicating that *Hivep3* deficiency does not trigger a compensatory upregulation of *Hivep1*. Because the mTEC maturation defect arose despite its presence, Hivep1 does not fully substitute for Hivep3, indicating a non-redundant requirement for Hivep3 in mTEC^High^.

**Figure 6:**
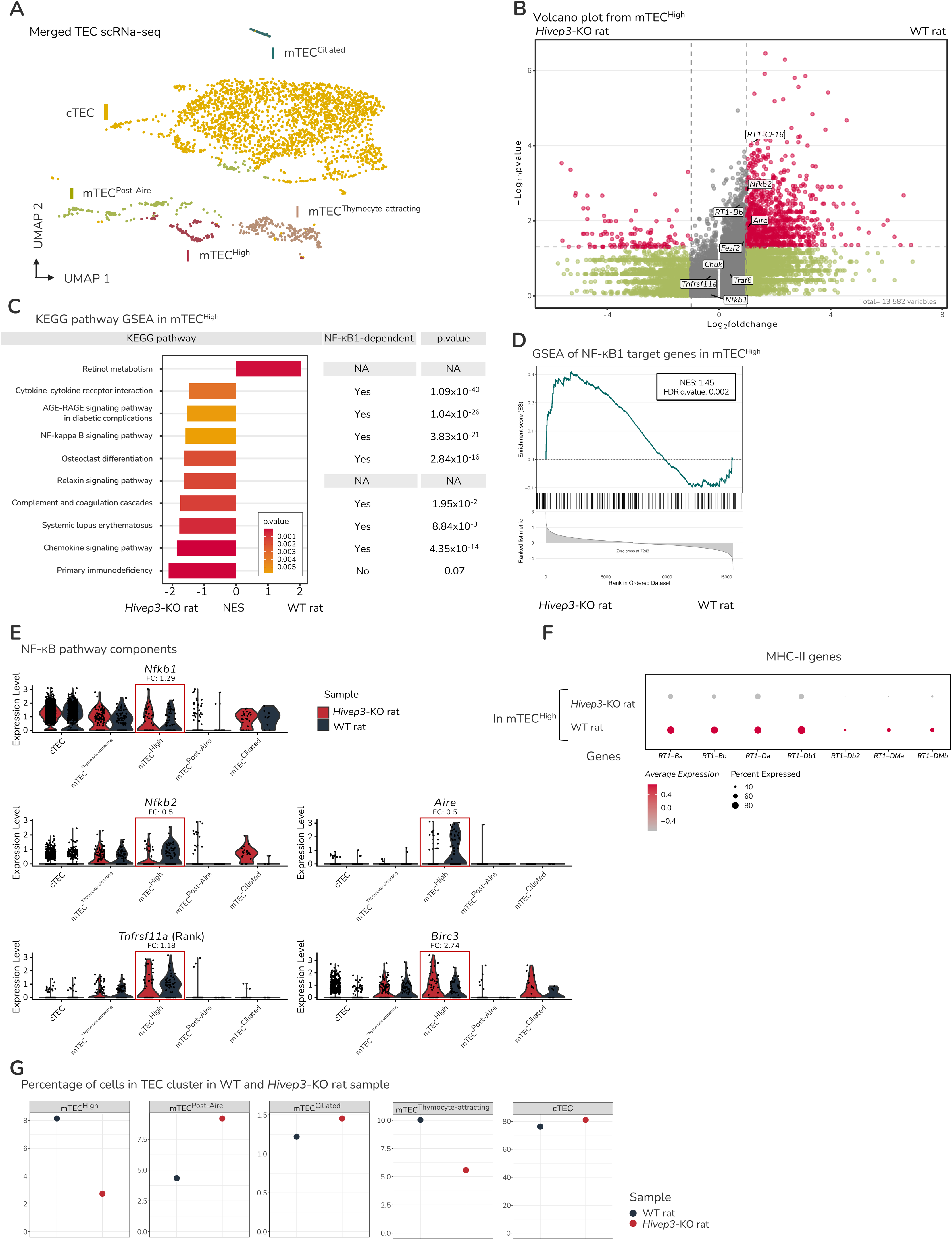
Hivep3 constrains canonical NF-κB1 and sustains non-canonical NF-κB2 signalling in. mTEC^High^ **(A)** Merged uniform manifold approximation and projection (UMAP) of scRNA-seq of *Hivep3*-KO and WT rat thymus. Each dot represents a cell. **(B)** Volcano plot of DE genes between *Hivep3*-KO and WT mTEC^High^. **(C)** Gene set enrichment analysis with KEGG pathways from DE genes between *Hivep3*-KO and WT mTEC^High^ showing enrichment of NF-κB1-associated pathways. **(D)** Gene set enrichment analysis of curated NF-κB1 target genes. **(E)** Violin plot of the expression of NF-κB pathway components across TEC subsets in *Hivep3*-KO and WT. **(F)** Dot plot showing expression of select MHC-II molecules in mTEC^High^ cells from *Hivep3*-KO and WT rats. **(G)** Proportions of thymic epithelial cell (TEC) subsets in *Hivep3*-KO and WT rat thymus.

We then performed a gene set enrichment analysis (GSEA) on KEGG pathways comparing *Hivep3*-KO and WT mTEC^High^. This showed that the majority of enriched pathways in *Hivep3-KO*, including cytokine-cytokine receptor interaction, AGE-RAGE signalling, chemokine signalling and complement and coagulation cascades, contained a significant over-representation of known NF-κB1 target genes (Figure 6C), indicating a broad activation of canonical NF-κB1-dependent transcriptional programs upon *Hivep3* deletion. This was further supported by the enrichment of the NF-κB signalling KEGG pathway itself, whose contributing genes were predominantly NF-κB1-dependent. Targeted GSEA using a curated NF-κB1 target gene set confirmed this canonical pathway activation in *Hivep3-KO* mTEC^High^ (NES = 1.45, FDR q = 0.002) (Figure 6D), despite mostly unchanged *Nfkb1* expression levels (Figure 6E). Among the genes contributing to these pathways, several pro-inflammatory mediators were upregulated in *Hivep3*-KO mTEC^High^, including the cytokines and receptors *Il6*, *Il6r* and *Tnfrsf18*, the alarmins *S100a8* and *S100a9*, and the complement components *C1qa*, *C1qb* and *C1qc*. Some of these genes, in particular *S100a8* and *S100a9*, can in turn activate canonical NF-κB through TLR4 and RAGE, a pattern-recognition receptor (PRR) for S100 alarmins, potentially amplifying the inflammatory program and further sustaining canonical NF-κB signalling activity in the KO. Consistent with a conserved repressive role, deletion of the paralog *Hivep1* increases the expression of NF-κB target genes and promotes systemic inflammation in mice^33^, which supports a conserved repressive function of HIVEP factors at κB-regulated inflammatory loci. This pattern is consistent with canonical NF-κB1 target activation arising from relieved competition at shared κB binding sites, given the overlapping DNA-binding specificity of HIVEP and NF-κB factors (Figures S1C, S2C), rather than from *Nfkb1* transcriptional upregulation, in line with the reported ability of HIVEP3 to bind the κB motif and inhibit NF-κB^20^.

Whereas *Nfkb1* expression remained largely unchanged, *Nfkb2*, encoding the central effector of the non-canonical NF-κB pathway, was significantly downregulated in *Hivep3*-KO mTEC^High^ (Figure 6B), indicating that Hivep3 is required to sustain *Nfkb2* expression in these cells. Of note, this reduction was unlikely to reflect downregulation of the receptor itself, as Rank (*Tnfrsf11a*) was expressed at comparable levels between genotypes (Figure 6E). Because non-canonical NF-κB signalling is essential at multiple stages of mTEC development and regulates *Aire* expression^34^, this reduced *Nfkb2* expression is consistent with Hivep3 sustaining mTEC maturation and Aire expression through the non-canonical NF-κB pathway. In addition to reduced *Nfkb2* expression, the activation of canonical NF-κB1 targets described above may further limit non-canonical signalling, as several of these targets converge on NIK, the central node of the pathway: Birc3 (cIAP2), upregulated in the KO (FC 2.74), directly promotes NIK degradation, while inflammatory amplifiers such as S100a8/9 (see above) sustain canonical NF-κB activity, itself a driver of NIK turnover. Both contributions are consistent with the canonical-to-non-canonical crosstalk reported in LTβR-stimulated stromal cells, where active canonical signalling and cIAP2 cooperate to destabilise NIK^35^. Together, these inputs could further limit p52/RelB activation, adding to the effect of reduced *Nfkb2* expression on non-canonical signalling in *Hivep3*-KO mTEC^High^.

The maturation defect of *Hivep3*-KO mTEC^High^ extended beyond *Aire* to MHC-II expression. All seven classical and non-classical MHC-II genes (*RT1-Ba*, *RT1-Bb*, *RT1-Da*, *RT1-Db1*, *RT1-Db2*, *RT1-DMa*, *RT1-DMb*) were reduced in *Hivep3*-KO mTEC^High^ (Figure 6F), four of them significantly (*RT1-Ba*, *RT1-Bb*, *RT1-Db1*, *RT1-DMa*; p < 0.05) and the remaining three trending in the same direction (p < 0.1) (Table 3). Concomitantly, the proportion of mTEC^High^ was reduced by approximately threefold in *Hivep3*-KO samples (Figure 6G), confirming that *Hivep3* deletion affects both the functional maturation and the abundance of mTEC^High^.

Collectively, these findings are consistent with a model in which Hivep3 exerts opposite effects on canonical and non-canonical NF-κB signalling. By competing for shared κB sites, Hivep3 constrains canonical NF-κB1 and limits the associated inflammatory program, while it sustains non-canonical NF-κB2 signalling, at least in part, by promoting *Nfkb2* expression. Because the activated canonical pathway in turn restrains NF-κB2 activity at the level of NIK, these effects converge to compromise mTEC maturation and *Aire* expression upon *Hivep3* deletion.

### Hivep3 promotes a distinct transcriptional program of TSA expression in mTEC^High^

Having found that Hivep3 sustains mTEC maturation, we next asked whether it also shapes the repertoire of TSAs expressed by mTEC^High^. To characterise Hivep3-induced genes in mTEC^High^, we selected those strongly upregulated in WT compared to *Hivep3*-KO samples (log₂FC > 2; 1,113 genes) and compared them with Hivep3-neutral genes (−1 < log₂FC < 1). To assess tissue specificity, we computed Tau scores from GTEx human RNA-seq^32^ for the Hivep3-induced genes that mapped to rat-human orthologs (845 out of 1,113). Hivep3-neutral genes were predominantly distributed at low Tau scores (< 0.5), indicating broad tissue expression, whereas Hivep3-induced genes shifted toward higher Tau scores, consistent with TSA expression (Figure 7A). Among the Hivep3-induced genes, TSA genes were predominantly associated with brain, testis, oesophagus, liver, retina, bone marrow and skeletal muscle (Figure 7B). To further quantify this bias, given the lack of analogous TSA classification in rats, we used the curated mouse TSA/non-TSA classification from *Sansom et al., 2014*^36^. After applying this classification, we observed a significant enrichment of TSAs among the Hivep3-induced gene set (60.6%; 321 genes) compared to the Hivep3-neutral set (34.2%; 183 genes; OR = 2.95, 95% CI 2.30–3.79; p < 1e-16), supporting a preferential induction of TSA genes by Hivep3 (Figure 7C). The remaining 583 Hivep3-induced and 578 Hivep3-neutral genes were excluded from this analysis, as they either lacked a clear classification in *Sansom et al., 2014*^36^ or had no rat-mouse ortholog. Next, to evaluate the Aire dependency of these 321 Hivep3-induced TSAs, we re-analysed the scRNA-seq data from heterozygous (HE) and *Aire*-KO rat TECs (*Stoljar et al., 2025* ^37^). We isolated the mTEC^High^ cluster and performed DE analysis between the two genotypes, allowing us to identify Aire-induced genes in the rat (Table 4). Strikingly, Aire-independent genes represented over three-quarters of Hivep3-induced TSAs, suggesting that Hivep3, although sharing a subset of targets with Aire, predominantly drives a distinct Aire-independent TSA program (Figure 7D). To gain insight into the mode of control of TSA expression by Hivep3, we assessed motif enrichment for HIVEP family TFs (Hivep1/3, Hivep2), NF-κB1 and NF-κB2 within the promoters of Hivep3-induced genes. Among these motifs, only the Hivep1/3 motif showed enrichment (odds ratio > 1.5) across all three categories, i.e., Hivep3-induced, Aire-independent TSAs and Aire-dependent TSAs (Figure 7E). This indicates that Hivep3 can induce TSA expression directly through its cognate recognition motif. Together, these data indicate that Hivep3 is able to induce TSA expression directly and indirectly through its effect on mTEC maturation.

**Figure 7:**
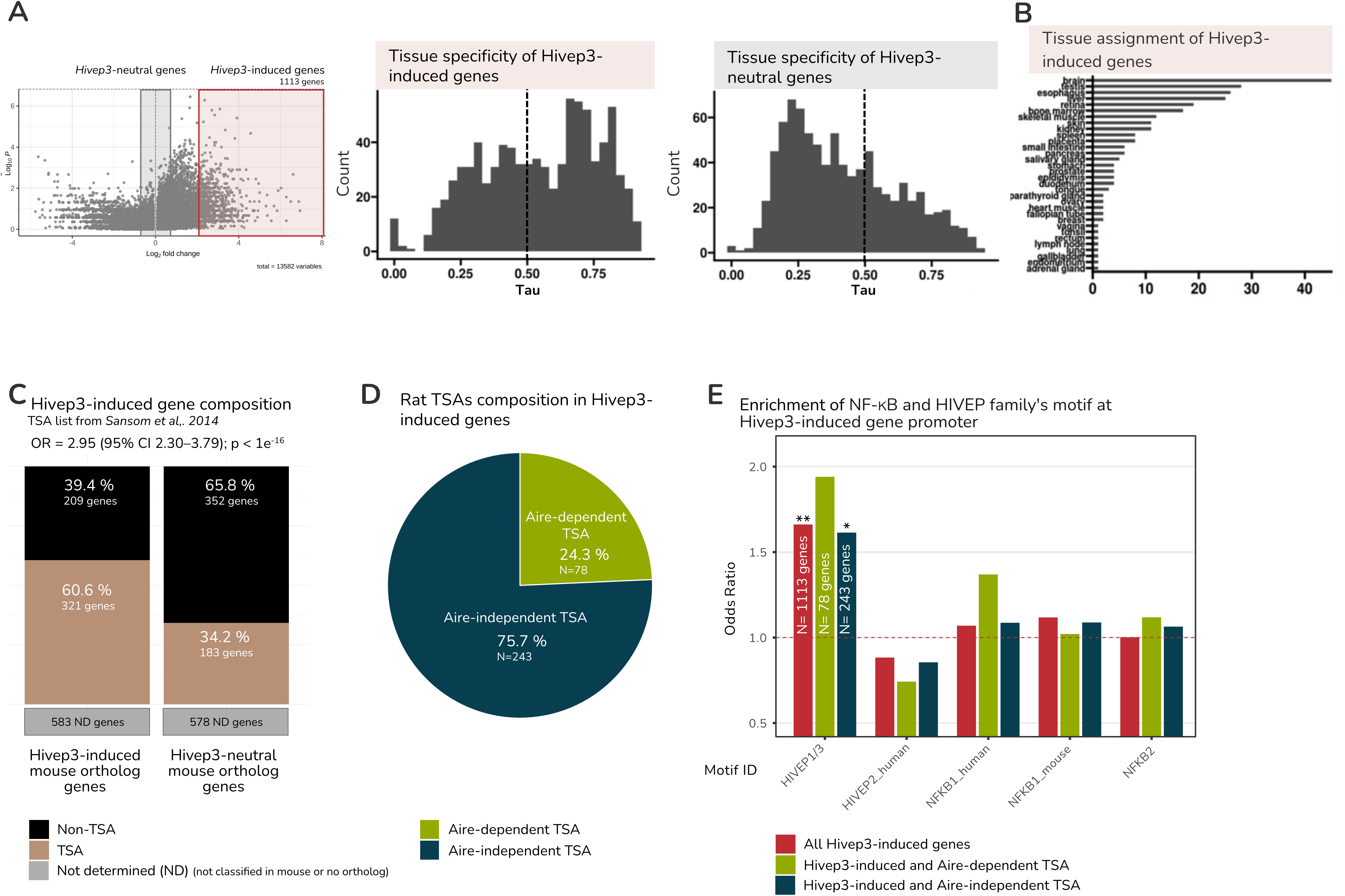
Hivep3 promotes a unique transcriptional program of TSA expression in mature mTECs. **(A)** Bar plot showing tissue specificity distribution of Hivep3-induced and Hivep3-neutral genes. **(B)** Bar plot illustrating tissue specificity of Hivep3-induced genes. **(C)** Stacked bar plot depicting the composition of Hivep3-induced versus Hivep3-neutral genes. The TSA gene list was obtained from rat orthologs of the mouse dataset from Sansom et al. (2014). Gene composition is colour-coded. **(D)** Pie chart showing the proportion of Aire-dependent and Aire-independent genes among Hivep3-induced TSAs (derived from mouse ortholog mapping). **(E)** Bar plot of motif enrichment at promoters of *Hivep3*-induced genes, colour-coded by category. (*p = 0.039; **p = 0.00074).

### Hivep3 shapes the cTEC transcriptional network and sustains efficient thymocyte positive selection

In rat scRNA-seq data, although *Hivep3* expression was highest in mTECs, it was also expressed in cTECs (Figure 4C). Consistent with a potential role in both TEC lineages, our scATAC-seq analyses revealed HIVEP motif enrichment in the human immature TEC cluster, compatible with bipotent progenitor identity (Figure 1E), and in mouse TAC-TECs, an early mTEC progenitor (Figure 2C-D). Within the human cTEC trajectory, this enrichment was confined to immature progenitors and was reduced in mature and transitional cTEC clusters (Table 1). To assess the impact of *Hivep3* deletion beyond the mTEC compartment, we performed DE analysis on cTECs from our WT and *Hivep3*-KO scRNA-seq data (Figure 8A, Table 3).

**Figure 8:**
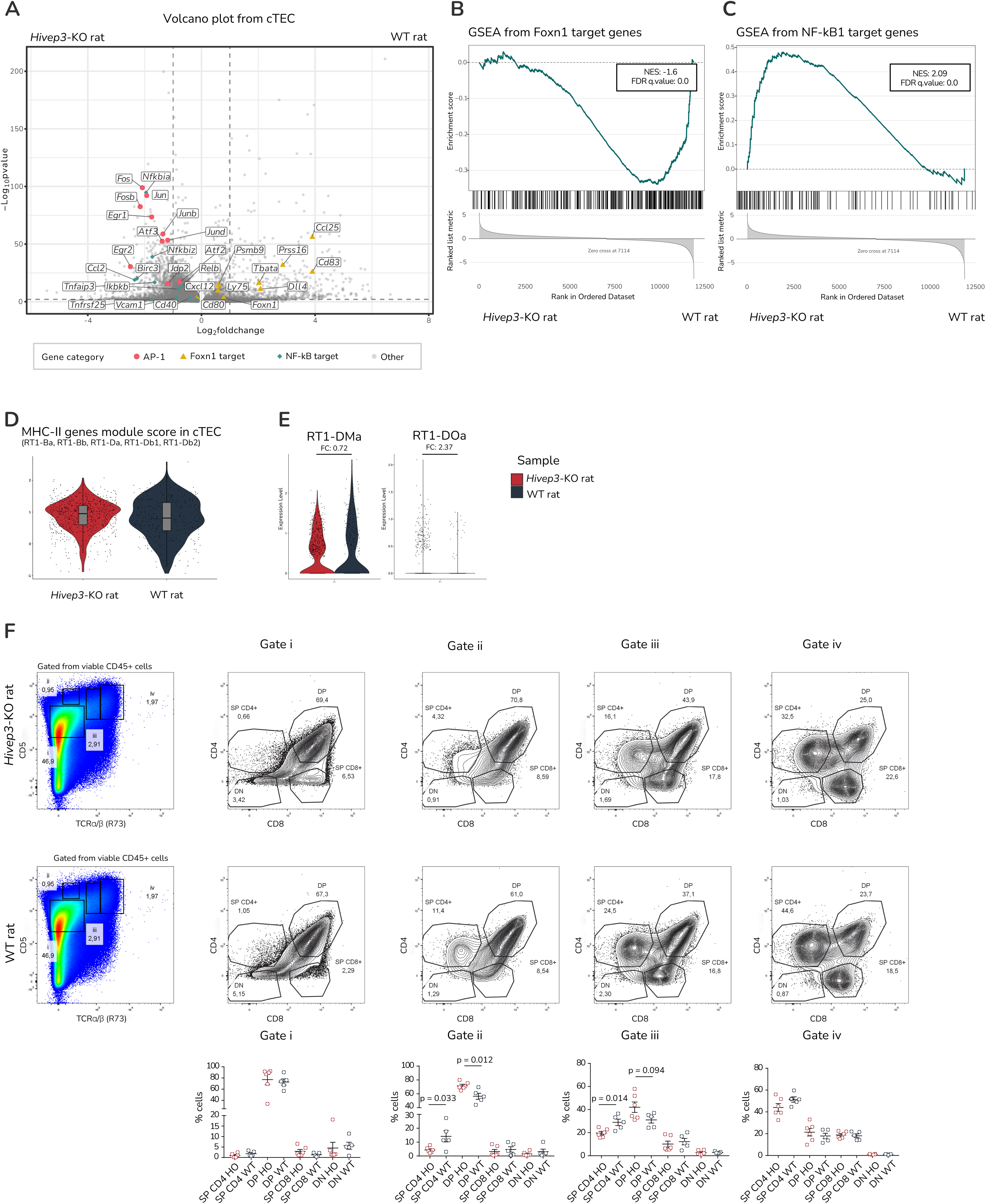
Loss of Hivep3 induces an imbalance in the cTEC transcriptional network and alters positive selection of thymocytes. **(A)** Volcano plot showing DE genes between *Hivep3*-KO and WT cTECs. Genes associated with AP-1 signalling are shown in pink, Foxn1 targets in yellow, NF-κB1 targets in blue, and all others in grey. **(B)** Gene set enrichment analysis of curated Foxn1, and **(C)** NF-κB1 target genes. **(D)** Violin plot depicting the module score of MHC-II-associated genes (*RT1-Ba*, *RT1-Bb, RT1-Da, RT1-Db1, RT1-Db2*) in cTECs. **(E)** Violin plot depicting the expression of non-classical MHC molecules *RT1-DMa* and *RT1-DOa* in cTECs. *Hivep3*-KO animals are shown in red, WT controls in blue. **(F)** Flow cytometry analysis of positive selection in the thymus, defined using a four-gate CD5/TCR strategy as previously reported^41^. Representative plots are shown for *Hivep3*-KO (top) and WT (middle) rats. Bar plot shows pooled analysis, with *Hivep3*-KO in red and WT in blue (bottom); each dot represents an individual animal.

The resulting volcano plot revealed divergent transcriptional signatures: Foxn1 target genes (e.g., *Prss16*, *Dll4*, and *Ccl25*) were downregulated in *Hivep3*-deficient cTECs, while NF-κB1 target genes and AP-1 family genes (e.g., *Fos*, *Jun, Junb*) were upregulated (Figure 8A). To confirm the opposing regulation of the Foxn1 and NF-κB1 programs, we performed GSEA using curated Foxn1 and Nfkb1 target gene sets. Foxn1 target genes were downregulated in *Hivep3*-KO cTECs (NES = –1.6, FDR q < 0.001), while NF-κB1 target genes were upregulated (NES = 2.09, FDR q < 0.001) (Figure 8B-C). As Hivep3 acts at κB sites, the concurrent AP-1 upregulation most likely accompanies NF-κB1 activation rather than reflecting a direct Hivep3 effect. Of note, NF-κB1 and AP-1 both require the limiting transcriptional co-factor CBP (Creb-binding protein), which is also a required co-factor for Foxn1-mediated transcription^38–40^. This raises the possibility that Hivep3, by constraining NF-κB1 and the accompanying AP-1 activity, preserves CBP availability for Foxn1-driven transcriptional programs. Interestingly, these pronounced perturbations did not substantially affect classical MHC-II expression in cTECs, as *RT1-Ba, RT1-Bb, RT1-Da, RT1-Db1*, and *RT1-Db2* remained comparable between genotypes (Figure 8D). In contrast, the non-classical MHC-II molecules controlling peptide editing were affected: RT1-DOa, the inhibitor of the RT1-DM editor, was markedly increased in *Hivep3*-KO cTECs (FC 2.37), while the editor RT1-DMa showed a minor reduction in the same direction (FC 0.72) (Figure 8E). Because RT1-DO restrains RT1-DM, both changes shift the DM/DO balance towards reduced peptide editing in *Hivep3*-KO cTECs, an effect driven primarily by the robust increase in RT1-DO. To determine whether these cTEC perturbations translate into altered thymocyte selection, we examined by flow cytometry the distribution of double negative (DN), double positive (DP) and single positive (SP) thymocytes across selection checkpoint gates, using a previously described four-gate CD5/TCR strategy^41^. In the gates capturing thymocytes undergoing positive selection (gates ii and iii), we observed a reduction of CD4⁺ SP thymocytes (significant in both gates) together with an accumulation of DP thymocytes (significant in gate ii; trend in gate iii, p = 0.094), with CD8⁺ SP thymocytes unaffected (Figure 8F). This CD4-selective impairment suggests that *Hivep3* deficiency compromises MHC-II-restricted positive selection or delays CD4⁺ thymocyte maturation, consistent with the altered DM/DO balance in *Hivep3*-deficient cTECs. To investigate whether the positive selection defect stems from a thymocyte-intrinsic requirement for Hivep3 or from cTEC-mediated dysregulation, we examined HIVEP family expression during rat thymopoiesis by scRNA-seq. Unlike *Hivep1* and *Hivep2*, *Hivep3* was selectively expressed in mature SP thymocytes poised for thymic egress (CCR7⁺CD69⁻) and remained low or undetectable in DN, DP and CD69⁺ thymocytes, encompassing the stages at which positive and negative selection take place (Figure S5C-D). This expression pattern argues against a thymocyte-intrinsic role for *Hivep3* in either positive or negative selection, and instead implicates a TEC-driven mechanism underlying the selection defect observed in *Hivep3*-KO rats. Together, these data indicate that Hivep3 fine-tunes the cTEC transcriptional network and that its deficiency disrupts the balance between Foxn1 and NF-κB1/AP-1 transcriptional programs, ultimately impairing CD4⁺ thymocyte positive selection.

### Loss of Hivep3 reshapes peripheral T-cell composition and differentiation

We next asked whether the thymic dysfunction observed in *Hivep3*-KO rats could lead to a systemic immune dysregulation. Therefore, we phenotyped main splenic immune populations, with primary focus on the T-cell compartment, as a representative readout of systemic immune homeostasis (Figures 9A-D).

**Figure 9:**
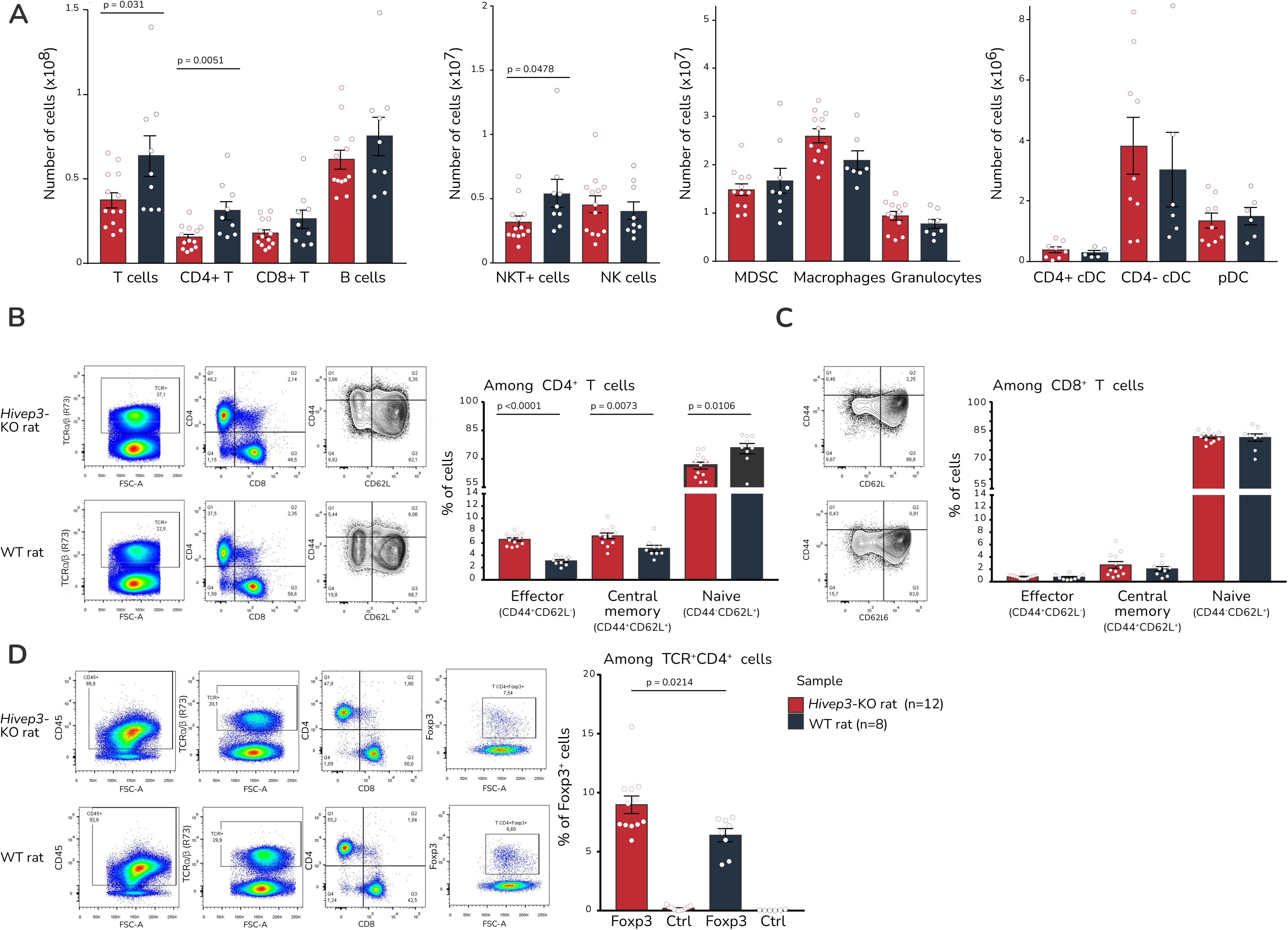
Hivep3 deficiency induces alterations in the immune repertoire. **(A)** Bar plot showing the number of splenic immune cells in *Hivep3*-KO (red) and WT (blue) rats. Flow cytometry gating strategy (right) and pooled data (left) showing the percentage of effector, memory, and naïve splenic **(B)** CD4⁺ T cells or (**C)** CD8⁺ T cells in *Hivep3*-KO (red) and WT (blue) animals. **(D)** Percentage of regulatory CD4⁺ T cells in *Hivep3*-KO (red) and WT (blue) animals.

First, in the spleen, independent of sex, we observed a significant reduction of the total T-cell numbers (p = 0.031), driven predominantly by CD4^+^ T-cells (p = 0.005), while CD8^+^ T-cells showed no significant difference (Figure 9A, Figure S6A). This observation parallels our thymic findings, where CD4^+^ T-cells were also preferentially affected by the deletion of *Hivep3* during thymic selection (Figure 8F). Among the other immune cells, while *Hivep3* deletion did not substantially impact B-cells, natural killer (NK) cells, dendritic cells (DCs), macrophages or granulocytes, we noted a significant reduction within the NKT^+^ cell population (p = 0.0478). This is consistent with previous reports that *Hivep3* deletion in mice reduces invariant natural killer T (iNKT) cells, a subset of the heterogeneous NKT^+^ population ^21,42^.

Within the splenic CD4^+^ T-cell compartment, further analysis revealed a significant shift in differentiation status. Indeed, the effector and central memory CD4^+^ T-cell (T_em_/T_cm_), subsets were significantly expanded (p < 0.01 for both), whereas naïve CD4^+^ T-cells were significantly reduced in *Hivep3*-KO rats (p = 0.0106; Figure 9B). In contrast, as observed previously, the CD8^+^ T-cell compartment remained unaffected (Figure 9C). We also examined the proportion of Foxp3^+^ Tregs within the CD4^+^ T-cell compartment and observed a significant increase in *Hivep3*-KO rats (p = 0.0214) (Figure 9D). These changes were independent of sex (Figure S6B). Next, to investigate whether Hivep3 loss exerts a cell-intrinsic effect on T-cell activation, we measured the proliferative response of WT and *Hivep3*-KO T-cells to polyclonal anti-CD3/CD28 stimulation *in vitro*. Across a range of plate-bound anti-CD3 coating concentrations (0-5 µg/mL), CD4^+^ T-cells from both genotypes showed comparable proliferative responses (two-way repeated-measures ANOVA, genotype effect p = 0.71). On the other hand, *Hivep3*-KO CD8^+^ T-cells exhibited a modest, non-significant increase in proliferation (Figure S6C). Theoretically, this CD8^+^ increase would be reflected by an expanded CD8^+^ population *in vivo*, which is the opposite of the reduction actually observed. Hence, the decrease of total and naïve CD4^+^ T-cells most likely arises from an altered thymic output, whereas the expansion of CD4^+^ T_em_ and T_cm_ cells reflects intact peripheral T-cell activation capacity in *Hivep3*-KO T-cells. Notably, this CD4-biased peripheral phenotype shares features with a previous report on *Hivep3* (then termed *Zas3*)-KO mice, in which *Allen et al., 2007* described an expanded T_em_ compartment and a reciprocal reduction of naïve cells within peripheral CD4⁺, but not CD8⁺, T-cells^43^. In that model, however, absolute peripheral CD4⁺ numbers were unchanged and the memory shift was ascribed to homeostatic proliferation, in contrast to the reduced CD4⁺ numbers and altered thymic output we describe above. Although the authors favoured a thymocyte-intrinsic role for *Hivep3* without direct evidence, our model therefore offers an alternative, TEC-centric reading of this CD4-biased shared peripheral phenotype.

### Loss of Hivep3 drives Th1-skewed systemic inflammation in rats

Next, given the expansion of the CD4^+^ T_em_ and T_cm_ cell compartment, we asked whether this was associated with a biased T helper response by conducting a longitudinal serum cytokine profiling of IFN-γ and IL-17A. *Hivep3*-KO rats displayed a sustained elevation of IFN-γ (p = 0.0404), a hallmark cytokine of Th1 responses, together with a decrease of IL-17A (p = 0.0004), characteristic of Th17 responses, across young (4-6W), adult (5-6M), and mature (8-9M) ages (Figure 10A). These cytokine alterations were similar in male and female rats (Figure S6D). Although iNKT-cells are a known cellular source of IFN-γ, their reduction in *Hivep3*-KO rats would by itself be expected to decrease, not elevate, iNKT-derived IFN-γ^44^. The elevated serum IFN-γ, therefore, most likely originates from the proportionally enriched splenic CD4^+^ T_em_/T_cm_ cell compartment, which produces more IFN-γ per cell than naïve T-cells and can sustain elevated cytokine levels despite the overall reduction in total CD4^+^ T-cell numbers. Together, this sustained Th1-skewed cytokine profile across age and sex indicates a chronic, CD4-driven Th1 polarisation in *Hivep3*-KO.

**Figure 10:**
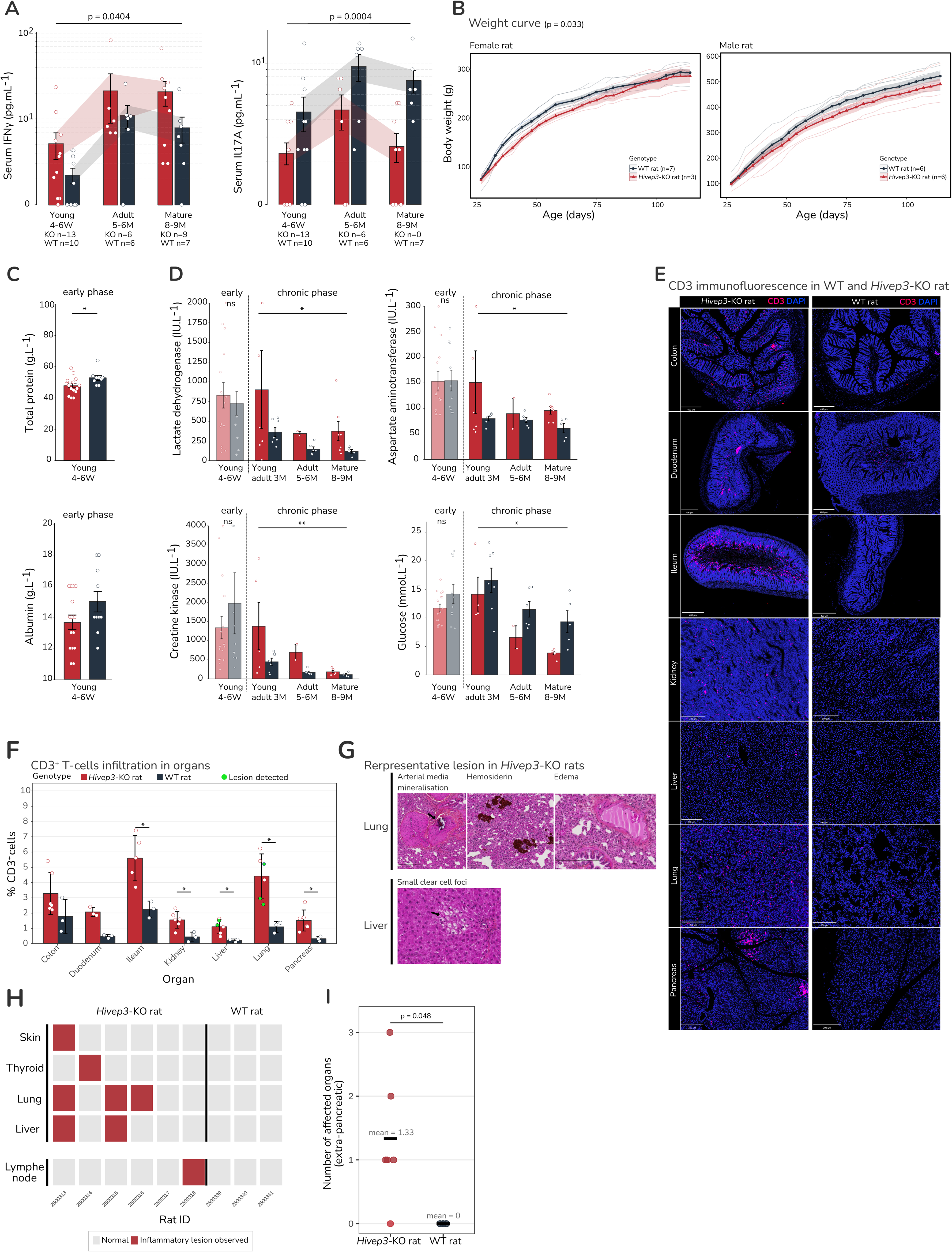
Hivep3 favours Th1-mediated chronic inflammation, and its loss induces inflammatory lesions. **(A)** Longitudinal analysis of IFNγ and IL-17A concentrations (pg·mL⁻¹) in the serum of young, adult, and mature *Hivep3*-KO (red) and control (blue) animals. **(B)** Weight curves of *Hivep3*-deficient (red) and WT (blue) rats from 27 to 114 days of age in female (left) and male (right) rats. p = 0.033. **(C)** Total protein (top) and albumin (bottom) concentrations in the blood of *Hivep3*-KO and WT rats. **(D)** Lactate dehydrogenase, aspartate aminotransferase, creatine kinase, and glucose concentrations in the blood of *Hivep3*-KO and WT rats. Two-way ANOVA was used. **(E)** CD3 immunofluorescence staining in *Hivep3*-KO (left) and WT (right) rats across colon, duodenum, ileum, kidney, liver, lung, and pancreas. CD3 in red and DAPI in blue. Scale is noted at the bottom. **(F)** Bar plot representing the percentage of CD3^+^ cells in colon, duodenum, ileum, kidney, liver, lung, and pancreas in *Hivep3*-KO (red) and WT (blue). A full circle represents a rat with observed inflammatory lesions. **(G)** Histological image of representative lesions in *Hivep3*-KO rats from lung and liver. **(H)** Plot showing the presence of inflammatory lesions in *Hivep3*-KO and WT rats across skin, thyroid, lymph nodes, lung, and liver. **(I)** Pooled plot showing the number of organs affected by inflammatory lesions in *Hivep3*-KO (red) and WT rats (blue). Each dot represents an individual animal.

This Th1 polarisation coincided with systemic perturbations, including a slowed body weight gain from weaning to adulthood (D24-D114; p = 0.033), independent of sex (Figure 10B). In addition, we noted hypoproteinaemia (p = 0.0263) and a trend toward lower albumin concentration in the early phase (4-6W), consistent with the well-documented catabolic effects of chronic IFN-γ activity^45^ (Figure 10C). From 3 months onward, these parameters normalised to levels comparable with WT controls, suggesting partial systemic adaptation to chronic inflammation, possibly aided by the expanded Treg compartment. Conversely, tissue injury markers showed no difference in the early phase but became significantly elevated from 3 months onward, including lactate dehydrogenase (LDH; p = 0.03) and aspartate aminotransferase (AST; p = 0.041), alongside a trend toward increased creatine kinase (CK; p = 0.083) in *Hivep3*-KO rats (Figure 10D). Together, this biochemical signature is consistent with ongoing multi-organ tissue injury, with elevated AST and LDH particularly suggestive of hepatic involvement, and the CK trend in line with the catabolic muscle protein turnover characteristic of chronic Th1-driven inflammation. Notably, hypoglycaemia (p = 0.004) was also observed in *Hivep3*-KO rats, consistent with the glycolytic demand of expanded CD4⁺ T_em_ and T_cm_ compartments and possibly compromised hepatic gluconeogenesis (Figure 10D).

Consistent with these systemic and biochemical alterations, in aged *Hivep3*-KO rats (≥1 year), tissue-level immunofluorescence revealed a clear accumulation of CD3^+^ T-cells across multiple organs, reaching statistical significance in the ileum, kidney, liver, lung, and pancreas (Figure 10E-F). Diverse lesions, including skin hyperkeratosis, lymph node T-cell area hyperplasia, minimal thyroid hyperplasia, focal pulmonary oedema with haemosiderin-laden macrophage accumulation and arterial media mineralisation, and multiple small clear hepatic foci, accompanied this widespread infiltration (Figure 10G). Overall, the histopathological scoring showed a significantly higher incidence of inflammatory lesions in *Hivep3*-KO animals, compared to controls (mean lesion score: 1.33 vs 0; p = 0.048; Figure 10H-I). This constellation is consistent with chronic T-cell-driven tissue pathology, in line with the sustained Th1 response established above.

Collectively, these findings establish Hivep3 as a critical regulator of immune homeostasis, whose deficiency compromises central tolerance and leads to a Th1-skewed inflammation, culminating in metabolic dysregulation and chronic multi-organ damage.

## DISCUSSION

Spanning paediatric human and mouse thymi, our study generated a cross-species chromatin atlas of mTECs to identify TFs underlying their maturation and TSA expression. We found that mTEC identity is governed by a dynamic TF landscape in which largely conserved core regulators are interwoven with species-specific features. Among these, alongside the well-known NF-κB family, HIVEP members emerged as conserved and prominent TFs with a key role in the regulation of TEC maturation. In mice, all three paralogs have been implicated in distinct facets of the immune system. Hivep1 has been reported to regulate Th17 differentiation in the context of nonalcoholic steatohepatitis (NASH)^46^. Hivep2 is both required for CD4 and CD8 positive selection through an intrinsic role in the haematopoietic compartment and restrains Th2 differentiation^47,48^. Hivep3 regulates iNKT differentiation and function, promotes IL-4 production in CD8^+^ iNKT-cells, and regulates the *IL2* gene in T-cells, while its deletion leads to increased bone formation^23,49,21,42^. In human T-cells, HIVEP2 has additionally been linked to regulatory T-cell function^50^. However, none of these studies addressed HIVEP function in TECs or in central tolerance.

Furthermore, these immune phenotypes were characterised primarily in mouse models, whereas rat models frequently diverge from mice in immune biology, often more closely recapitulating human physiology^51–54^. Here, the strong and conserved accessibility of HIVEP recognition motifs in immature and *AIRE*-accessible mTECs implicated the HIVEP family in mTEC regulation, although the motif cannot discriminate between paralogs. Among the co-expressed paralogs *Hivep3* and *Hivep1*, we focused on *Hivep3*.

Therefore, we generated a CRISPR/GONAD-mediated *Hivep3*-KO rat to interrogate the role of Hivep3 in thymic epithelial biology and central tolerance. In the thymus, we found that *Hivep3* deficiency led to medullary islet fragmentation, expansion of the Krt14^+^ compartment, and a marked reduction in Aire^+^ cells, indicating impaired mTEC maturation and potentially compromised TSA expression. At the transcriptomic level, deletion of Hivep3 induced opposite effects on canonical and non-canonical NF-κB signalling: *Nfkb2* expression was significantly reduced, whereas the NF-κB1-driven pathway and its targets were increased despite unchanged *Nfkb1* transcript levels. This canonical activation, occurring without a change in *Nfkb1* expression, is consistent with a previous study in *Zas3*(*Hivep3*)-KO mouse tissues, where its deletion increased nuclear RelA/p50 DNA binding across spleen, brain and liver by EMSA^43^. Complementing these data, an ATAC-seq study in mouse iNKT-cells showed that κB-binding motifs were more accessible in *Hivep3*-deficient cells than in control^21^. In this framework, Hivep3 appears to constrain canonical NF-κB1-driven transcriptional programs through competitive occupancy of κB binding sites, while simultaneously sustaining non-canonical NF-κB2 signalling, at least in part, by promoting *Nfkb2* expression. Mechanistically, our findings suggest that these two NF-κB pathways are linked rather than independently regulated. Upregulation of the canonical NF-κB1 targets, including negative regulators of NIK, such as *Birc3*, provides a plausible basis for the attenuation of p52/RelB activity, which is consistent with established canonical to non-canonical NF-κB crosstalk. An alternative, non-mutually exclusive possibility is that Hivep3 may directly regulate components of the canonical and non-canonical pathway. Indeed, a study in RAW264.7 macrophages demonstrated that Hivep3 directly binds TRAF6, enhances its expression, and increases RelB expression upon RANKL stimulation^55^. Whether such interactions also occur in primary TECs, and whether Hivep3 directly engages with other regulatory elements of the NF-κB pathway, remain to be determined. Addressing these questions is currently constrained by limited rat-specific reagents, notably a validated anti-Hivep3 antibody, and by the low yield of primary mTEC^High^ from rat thymus. Nevertheless, our findings support a model in which Hivep3 functions as a regulatory switch, balancing the NF-κB pathway by linking κB motif competition with pathway crosstalk to control Aire and mTEC maturation.

In addition to its effect on mTEC lineage progression, Hivep3 also promotes a distinct TSA transcriptional program. Notably, over three-quarters of Hivep3-induced TSAs were Aire-independent, defining Hivep3 as a regulator of an Aire-independent TSA axis and a previously unrecognised major player of central tolerance. As a caveat, although our Aire-dependency analysis was performed rat-on-rat, the TSA classification itself relied on rat-to-mouse ortholog mapping in the absence of a curated rat TSA reference. The precise breadth of Hivep3-regulated TSAs will require refinement once such a resource becomes available.

In parallel, in cTECs, deletion of *Hivep3* disrupts the NF-κB1 transcriptional network, and alongside it, the Foxn1-driven transcriptional program. Foxn1 is a master regulator of TEC identity and function, yet the factors that fine-tune its activity *in vivo* remain unresolved. Here, *Hivep3* deficiency compromised the Foxn1 program in cTECs, revealing a previously unrecognised requirement for *Hivep3* in sustaining Foxn1 output in this lineage. At the chromatin level, our scATAC-seq data indicate that HIVEP motifs are accessible in the human Immature TEC cluster (7^th^), a bipotent progenitor of both cTEC and mTEC lineages, but reduced in mature or transitional cTEC clusters (101^st^ and 479^th^, respectively). This raises the possibility that the transcriptional alteration observed in cTECs, or at least part of it, originates from earlier perturbations in Immature TEC, with mature cTECs inheriting an imbalance established at the progenitor stage, where *HIVEP3* is normally active. In addition, given that *Hivep3* is expressed in WT rat cTECs, a direct cell-intrinsic contribution cannot be excluded, possibly through protein-level interactions of Hivep3 with the NF-κB1 or Foxn1 machinery. In this model, developmental and cell-intrinsic mechanisms may cooperate to reshape the cTEC transcriptional landscape. Taken together, because NF-κB1, AP-1 and Foxn1 all depend on the limiting co-activator CBP/p300, we hypothesise that the increased NF-κB1 and AP-1 activity in *Hivep3*-deficient cTECs competes for CBP/p300 at the expense of Foxn1, a phenomenon known as “CBP squelching”, whereby limited co-activator availability reshapes transcriptional output. Functionally, we observed that *Hivep3* expression was restricted to post-selection CCR7⁺CD69⁻ SP thymocytes, and absent from earlier DN, DP and CD69^+^ stages where selection occurs. Hence, the reduced CD4^+^ output cannot reflect a thymocyte-intrinsic requirement, but instead supports a stromal contribution, specifically from cTECs.

In the periphery, these thymic alterations manifest as systemic, Th1-skewed inflammation, marked by an expansion of CD4^+^ effector and central memory T-cells (T_em_/T_cm_), an increased proportion of Foxp3^+^ Tregs within the CD4^+^ compartment, elevated serum IFN-γ, and reduced IL-17A, and marked CD3^+^ T-cell infiltration across multiple organs, accompanied by inflammatory lesions. Importantly, *in vitro* anti-CD3/CD28 stimulation revealed comparable proliferative responses in *Hivep3*-KO and WT T-cells, further excluding a T-cell-intrinsic activation defect and supporting that the peripheral phenotype reflects altered thymic output rather than a cell-autonomous role of Hivep3 in T-cells. This interpretation also reconciles our findings with the two earlier murine *Hivep3* models, neither of which established a thymocyte-intrinsic basis for the comparable CD4-biased phenotype. *Allen et al., 2007* reported a reduced thymic CD4/CD8 ratio and an expansion of memory CD4⁺ cells in *Zas3*-deficient mice. Yet peripheral T-cell numbers were unchanged, and the authors attributed the memory shift to homeostatic proliferation following reduced CD4⁺ thymic output. The reduction in absolute splenic CD4⁺ numbers in our model, by contrast, indicates an output deficit rather than peripheral conversion, and their study included neither stage-resolved expression nor TEC analysis to support a cell-intrinsic role. In contrast, a strictly thymocyte-intrinsic requirement in selection has been well-established for the paralog *Hivep2* by reciprocal bone marrow chimaeras, in which *Hivep2*-deficient thymic epithelium supported normal T-cell development, whereas *Hivep2*-deficient bone marrow did not^47^. This thymocyte-intrinsic requirement, together with the uniform expression of *Hivep2* across thymocyte stages, contrasts with the post-selection, SP-restricted expression of *Hivep3* in our model. Thus, although a thymocyte-intrinsic contribution cannot be formally excluded, our data collectively support a stromal, TEC-centred mechanism for *Hivep3* in rats.

These observations raise a central question: given that *Hivep3* deficiency disrupts cTEC function, impairs mTEC maturation, and reduces TSA expression, why does the resulting peripheral phenotype manifest as Th1-biased inflammation rather than fulminant autoimmunity? We propose two non-mutually exclusive mechanisms.

First, one possible explanation is the functional compensation within the HIVEP transcription factor family, which may buffer the full impact of *Hivep3* deficiency. The *Stoljar et al., 2025* data had shown a predominant *Hivep3* expression in rat TECs compared with other paralogs, which initially led us to expect a reduced likelihood of paralog compensation. Unexpectedly, our subsequent scRNA-seq analysis in the WT rat TECs revealed comparable expression between *Hivep1* and *Hivep3*, mirroring the pattern in the mouse. Thus, the co-expression we sought to avoid is also present in rats. Notably, the co-expression of both paralogs in rodent TECs suggests an evolutionary retention to potentially cooperatively regulate the κB-dependent transcriptional programs in the thymic stroma. However, in mTEC^High^, deletion of *Hivep3* did not lead to an upregulation of *Hivep1*, suggesting the absence of transcriptional cross-regulation between the two paralogs. These findings indicate that the full extent of HIVEP function in thymic epithelial biology may only be revealed through combinatorial perturbation, such as dual Hivep1/Hivep3 deficiency or complete ablation. However, given the broad developmental roles of HIVEP proteins, complete ablation of multiple paralogs, particularly all three, risks disrupting essential developmental programs, potentially leading to perinatal lethality or skeletal defects, which would confound interpretation of any thymic phenotype. To circumvent these caveats, future studies employing TEC-specific combinatorial genetic perturbations, such as Foxn1-Cre-driven deletion of floxed HIVEP alleles, may be used to disentangle paralog-specific and redundant functions *in vivo*. Complementary *ex vivo* approaches, such as CRISPR-dCas9-mediated repression (KRAB) or activation (VP64) in primary TEC cultures, may further refine the dissection of Hivep-dependent regulatory networks.

Second, the expansion of Foxp3^+^ Tregs may counterbalance the escape of autoreactive T-cells. Although the reduction of Aire^+^ cells and impaired TSA expression would be expected to generate self-reactive clones, the peripheral phenotype is not characterised by overt tissue-specific autoimmunity. Instead, we propose that the expanded Treg compartment restrains the effector function of these T_em_/T_cm_ populations, thereby limiting autoreactivity. In this model, *Hivep3* deficiency creates a “leaky” central tolerance checkpoint that permits the export of self-reactive clones, but concomitant Treg expansion contains their pathogenic potential. In addition to this regulatory context, the cytokine milieu is skewed towards Th1 immunity, with reduced IL-17A and elevated IFN-γ. This profile is consistent with the expanded T_em_/T_cm_ pool that arises from altered thymic output, rather than with iNKT-derived cytokines, which are reduced in *Hivep3*-KO rats and would otherwise predict the opposite IFN-γ trend. This selective bias is notable, given that several autoimmune diseases, such as rheumatoid arthritis and multiple sclerosis, involve both Th1 and Th17 activation^56^. One possible explanation is the preferential suppression of Th17 responses by the expanded Treg compartment, as reported in specific Treg subsets, thereby limiting a Th17-driven autoimmune phenotype^57,58^. In this context, while IFN-γ–driven inflammation is present, the lack of Th17-derived IL-17 may limit the transition from subclinical inflammation to overt autoimmunity. Consistent with this model, the immune imbalance observed in *Hivep3*-deficient animals manifests as chronic, low-grade inflammation accompanied by early catabolism, hypoglycaemia, late tissue damage, and progressive multi-organ inflammation, most likely driven by infiltrating CD4^+^ T-cells that originate from defective thymic output.

Collectively, these findings not only identify Hivep3 as a previously unrecognised regulator of TEC function and central tolerance whose deficiency leads to systemic Th1-biased inflammation, but they also suggest broader implications for human disease. In future studies, it will be important to determine whether HIVEP3 mutations or polymorphisms are associated with disorders characterised by Th1-biased signatures. More broadly, our findings illustrate a conceptual framework in which impaired central tolerance can drive peripheral T-cell expansion accompanied by compensatory Treg responses, without necessarily progressing to fulminant autoimmune tissue destruction. As exemplified by *Hivep3* deficiency in our model, this concept may help the interpretation of immune-related GWAS loci that do not map cleanly onto classical autoimmune phenotypes.

## METHODS

### Human samples

Human thymic samples were obtained as discarded and deidentified surgical waste from paediatric cardiac surgery performed at CHU de Nantes. All procedures were conducted in compliance with the French CODECOH regulatory framework under institutional declaration DC-2017-2987. A written agreement was established between CHU Nantes, the clinical team, and the research teams. Samples were stored in RPMI and fully anonymised before transfer to the research team, with only donor sex and date of birth retained.

### Isolation of paediatric human mTEC

Human thymic samples were obtained as deidentified samples from donors undergoing paediatric cardiothoracic corrective surgery at CHU Nantes. Samples were collected from both male and female donors, aged up to 2 weeks. Thymic tissue was vigorously minced with scissors and washed several times with cold RPMI. The lymphocyte-rich supernatant was discarded, and the remaining tissue fragments were enzymatically digested at 37°C for 35 min using an RPMI solution containing Collagenase D (Roche, #37334224), Dispase II (Sigma-Aldrich, #D4693), and DNase I (Sigma-Aldrich, #DN25). Following digestion, CD45⁺ haematopoietic cells were depleted by magnetic separation. The remaining cells were stained with EpCAM-PE (Miltenyi, clone HEA-125, #130-113-264) and positively selected using MojoSort™ Human anti-PE Nanobeads (BioLegend, #480092). For flow cytometric sorting, cells were stained with EpCAM-PE, CD205-APC (BioLegend, clone HD30, #342208), HLA-DR–PerCP-Cy5.5 (BD, clone G46-6, #560652), and DAPI for viability. Cell sorting was performed on a BD FACSAria instrument to isolate viable EpCAM^+^, CD205^-^and HLA-DR^+^ cells.

### Isolation of mouse mTEC

Thymi were collected from 4-week-old wild-type female C57BL/6 mice (Charles River) and finely minced with scissors to release thymocytes. Tissue fragments were then digested both mechanically and enzymatically. The first digestion step was performed using Collagenase D (Roche, #37334224) and DNase I (Sigma-Aldrich, #DN25) for 10 min at 37°C. This was followed by a second digestion using Dispase II (Sigma-Aldrich, #D4693) and DNase I for 30 min at 37°C. The resulting cell suspension was filtered and subjected to magnetic-activated cell sorting according to the manufacturer’s instructions. Enrichment was carried out on autoMACS (Miltenyi Biotec) using CD45 MicroBeads, mouse (Miltenyi, #130-052-301), to deplete haematopoietic cells, followed by positive selection using CD326 (EpCAM) MicroBeads, mouse (Miltenyi, #130-105-958) to isolate epithelial cells.

### Human and mouse scATAC-seq profiling

Nuclei isolation was performed on freshly isolated primary cells following the 10x Genomics Single Cell ATAC protocol (Documents CG000169 and CG000496). Sample processing, library preparation and sequencing were performed using the 10x Genomics Chromium platform and recommended scATAC-seq parameters (https://www.10xgenomics.com/support/epi-atac/documentation).

### Human and mouse scATAC-seq data processing

Data processing was performed using Cell Ranger ATAC (v2.1.0) using the hg38 genome for human and Cell Ranger ATAC (v1.2.0) using the mm10 genome for mouse. Downstream analysis of all scATAC-seq datasets was performed using the ArchR package (Granja et al., 2021^59^, v1.0.3), with default parameters unless stated otherwise. Arrow files were generated after filtering cells using the following criteria: for human files, *filterTSS = 8* and *filterFrags = 10,000* were applied, whereas for mouse, *filterTSS = 2* and *filterFrags = 2,000* were used. In both analyses, fragments mapping to mitochondrial DNA and sex chromosomes (X and Y) were excluded to avoid sex-based clustering artefacts and doublets were removed using *filterDoublets()*. For human samples, we applied Harmony batch correction and retained cells exhibiting a closed PTPRC locus. Peak calling was performed using MACS2. scRNA-seq integration, gene activity score calculation, motif and feature enrichment, chromVAR deviation analysis and trajectory analysis were performed following the user’s guide to ArchR (https://www.archrproject.com/bookdown/index.html). The Catalogue of Inferred Sequence Binding Preferences (CIS-BP) database, using either the Mus musculus or Homo sapiens reference, was used for TF motif enrichment analyses.

### Mouse scRNA-seq profiling

Thymic cell suspension from 6-week-old C57BL/6 mice was enriched for TECs by first magnetically depleting CD45^+^ cells using CD45 MicroBeads, mouse (Miltenyi, #130-052-301), before sorting EpCAM^+^UEA-1^+^Ly51^low^ cells using a FACS Aria III cell sorter (BD Bioscience). The thymic suspension was then prepared for single-cell sequencing, following 10x Genomics instructions. The libraries were prepared using the Chromium Single Cell 5’ v2 kit, with dual indexes, before being sequenced on an Illumina NextSeq 2000.

Data processing was performed using CellRanger (v6.1.2), and the reads were aligned to the mouse genome (mm10) with the --include-introns option. The downstream and QC analysis were performed using standard Seurat workflows (v4) by filtering out cells with *500 < nFeature_RNA < 6,000* and *mitochondrial transcript content > 6%*.

### Generation of *Hivep3*-KO Sprague-Dawley rat model and genotyping

All animal experiments were approved by the Animal Experimentation Ethics Committee of the Pays de la Loire region (France) under the permit number Apafis 26567, following the guidelines from the French National Research Council for the Care and Use of Laboratory Animals. All procedures were conducted in compliance with the European Community guidelines for laboratory animal care and use, and every effort was made to minimise animal numbers and their suffering.

The rat line carrying a deletion in Hivep3’s exon 1 was generated using CRISPR-Cas9 technology. A single-guide RNA (sgRNA) targeting 328 bp within the first exon was designed, produced, and validated by the TRIP platform (Nantes, France). To ensure specificity and efficiency, the sgRNA sequence was selected using CRISPOR, ChopChop, and IDT. The following sequences were used: sg-rHIVEP3-1191 R: 5’-ACGATAGTGACTCCCGGTAC-3’ ssODN:5’GCCCAGGAGGAGTTCCCTAACCAGGCGCAGCAGCGTGGAGTCCCCAAAATCCAGtCT cTAgaGatgaTCcCTcTCcTCCCACGGGGAGAAGACAAAACAGGAACAGTCACTGCTGAGCCTCCAGC-3’ sgRNA was transcribed *in vitro* with the HiScribe T7 high yield RNA Synthesis kit (New England Biolabs, Evry, France) and purified using EZNA microelute (Omega Bio-Tek, VWR, Rosny sous bois, France). Ribonucleoprotein (RNP) complexes were assembled by incubating sgRNA (0.2 µg/µL) and Cas9 protein (3 µM) at room temperature for 10 min, then maintained at 4°C prior to delivery using the GONAD method.

For genotyping, ear biopsies were collected from 10-day-old rats. Samples were digested overnight at 56°C in lysis buffer, diluted 1:20 in ultrapure water, and added to a PCR mix reaction prepared with Herculase II Fusion DNA Polymerase (Agilent Technologies), according to the manufacturer’s protocol. PCRs were carried out using the following primers:

Forward primer 5’- AGATCATCTTCGGCAAGTGTGG-3’, Reverse primer 5’- ACTGTACTCAGCTCCCAAAGGT-3’. Amplification was performed on a Veriti Thermal Cycler (Applied Biosystems) under the following conditions: 95°C for 5 min, 62°C for 2 min, 35 cycles of 95°C for 10 sec, 60°C for 10 sec, and 72°C for 30 sec, and a final extension at 72°C for 3 min. To assess mutations, PCR product lengths were analysed by heteroduplex mobility assay using a microfluidic capillary electrophoresis system (Caliper LabChip GX, PerkinElmer).

### Immunofluorescence staining

Thymic tissue was fixed in PFA, washed in PBS, snap frozen in OCT (Microm Microtech, #TFM-5), and frozen on dry ice before storage at-80°C. Cryosections of 6-7 µm were cut at-20°C onto SuperFrost Plus Adhesion microscope slides (Epredia, #10149870), air-dried overnight before storing at-20°C.

Frozen sections were fixed in 4% PFA in PBS (15 min, RT; Electron Microscopy Sciences, #15714-S), permeabilised with 0.1% Triton X-100 in PBS (15 min, RT; Sigma, #9002-93-1), and incubated with blocking buffer 10% NDS (Sigma-Aldrich, #D9663), 1% BSA, 0.01% Triton X-100, PBS (60 min, RT) to reduce nonspecific binding. Primary antibody staining was performed overnight at 4°C using Cytokeratin 14 (Novus Biologicals, #NBP2-89198) diluted in 1% BSA, 0.01% Triton X-100, PBS, followed by goat anti-rabbit AF647 secondary antibody (60 min, RT; ThermoFisher Scientific, #A-21245). Anti-AIRE-AF488 antibody (ThermoFisher Scientific, #53-5934-82) was then incubated for 120 min at RT, and DAPI (ThermoFisher Scientific, #62248) was used for nuclear counterstain. Slides were mounted with ProLong Gold Antifade Mountant (Invitrogen, P36934). Images were acquired on a Nikon Confocal A1R and a Hamamatsu NanoZoomer HT2 slide scanner.

### WT and *Hivep3*-KO rat scRNA-seq preparation

For single-cell suspension, thymi from 8–9-week-old WT and *Hivep3*-KO rats were collected, with two thymi pooled per genotype, finely minced with scissors to release thymocytes, washed 3 times in RPMI, and digested with Collagenase D (Roche, #37334224), Dispase II (Sigma-Aldrich, #D4693), and DNase I (Sigma-Aldrich, #DN25) for 40 min at 37°C. The resulting cell suspension was filtered and enriched for thymic epithelial cells by magnetic CD45 depletion and FACS sorting using CD45^-^ (BD, clone Ox-1, #561586), Epcam^+^ (Abcam, clone GZ-1, #Ab187276), anti-mouse IgG1 (BD, clone 10.9, #7432259), RT1-Bu^+^ (CR2TI, Ox3), and FVD506 (BD, #65-0866-14) using a BD FACSAria. The thymic suspension was then prepared for single-cell sequencing, following 10x Genomics instructions. The libraries were prepared using the Chromium Single Cell 3’ v3 kit and subsequently sequenced.

Data processing was performed using CellRanger (v6.1.2), and the reads were aligned to the custom rat reference based on the mRatBN7.2 genome with the --include-introns option. The downstream and QC analysis were performed using standard Seurat workflows (v5.3.0). Briefly, cells were filtered according to the following criteria: *nCount_RNA ≥ 500, nFeature_RNA ≥ 350, log10(nFeature_RNA) / log10(nCount_RNA) > 0.80*, and *mitoRatio < 0.20*. We then performed integration of the WT and *Hivep3-KO* datasets using the standard Seurat scRNA-seq integration framework (https://satijalab.org/seurat/articles/integration_introduction.html). After clustering, TEC clusters were subset using *Ptprc* and *Epcam* transcript levels, before constructing the neighbour graph using the first 30 dimensions in *FindNeighbors()* and setting the resolution to 0.4 using *FindClusters().* Cell populations were identified based on canonical marker genes. DE analysis between WT and *Hivep3*-KO conditions was performed at the single-cell level using Seurat’s *FindMarkers()* function with default parameters. Log2 fold change (avg_log2FC) values were used for downstream analyses.

### KEGG and transcription factor (NF-κB1 and Foxn1) gene set enrichment analysis

Gene set enrichment analysis was performed using the clusterProfiler package (v4.10.1). Differentially expressed genes were ranked by avg_log2FC. Gene symbols were converted to Entrez IDs using the org.Rn.eg.db annotation package (v3.18.0). A ranked gene list was generated and used as input for KEGG pathway enrichment analysis using the *gseKEGG()* function with the following parameters: *organism = “rno”, minGSSize = 5, maxGSSize = 500, pAdjustMethod = “BH”, and pvalueCutoff = 1*. Non-informative entries were removed prior to interpretation.

For NF-κB1, target genes were obtained from the TRRUST database^60^ (mouse dataset) while Foxn1 target genes were compiled from the mouse dataset reported by Žuklys et al. (2016)^61^. Gene set enrichment analysis was performed using the GSEAPreranked module in GenePattern (Broad Institute, v7.4.0) on a ranked list of DE genes with no_collapse, weighted scoring, meandiv normalisation, and 1,000 permutations.

### Hivep3-induced genes composition

To assess the composition of the *Hivep3*-induced genes in mTEC^High^ clusters, DE genes were first stratified based on log2 fold change, and filtered using |avg_log2FC| ≥ 2, while Hivep3-neutral genes were defined as those with avg_log2FC close to zero.

As no rat-specific TSA gene set is available yet, we used a previously published mouse TSA gene set from Sansom et al. (2014). *Hivep3*-induced or-neutral genes were classified as follows: TSA genes (mouse ortholog), non-TSA genes (mouse ortholog), and not determined (ND) due to the absence of ortholog mapping or absence from the original mouse TSA gene list.

To evaluate the Aire-dependence of these mouse orthologs *of Hivep3*-induced or-neutral TSA genes, we generated an Aire-dependent gene list using rat scRNA-seq data from Stoljar et al. (2025). Pseudobulk counts were generated by summing raw UMI counts per gene across all cells within each sample using Seurat. DE analysis between the samples was performed using DESeq2 (v1.50.2). Aire-dependent genes were defined as genes with log2FC ≤ −2. Rat TSA genes (derived from mouse ortholog mapping) were intersected with the list of Aire-dependent genes to determine the proportion of TSA genes regulated by Aire.

### Gene tissue specificity calculation and TSA enrichment analysis

Gene tissue specificity score and TSA enrichment analysis were calculated using the GeneTissueSpec Shiny application developed by Provin et al. (2026), available at: https://nathanprovin.shinyapps.io/GeneTissueSpe/. Tissue specificity index (Tau) was calculated for human ortholog genes using GTEx version 8 tissue expression data. Tau values range from 0 to 1, where values close to 1 indicate tissue-specific expression and values close to 0 indicate ubiquitous expression across tissues. TSA enrichment analysis was performed on the 1,113 Hivep3-induced genes defined by avg_log2FC ≥ 2.

### Body weight monitoring

Rats in each group were weighed at 11:00 AM on days 27 to 114, twice weekly. The study included 12 males (WT n = 6 and *Hivep3*-KO n = 6) and 10 females (WT n = 7 and *Hivep3*-KO n = 3). Statistical tests were performed using a linear mixed model (LMM) following the model: weight ∼ Day * genotype * sex + (1|rat_id).

### Serum biochemical measurements

Blood samples were collected and centrifuged at 3,000 × g for 15 min to obtain serum. The concentrations in serum of total protein, albumin, lactate dehydrogenase, aspartate aminotransferase, creatine kinase, and glucose were measured by Laboratoire de Biologie Médicale (CHU de Nantes, Nantes, France). In the early phase, we tested 10 WT and 16 *Hivep3*-KO rats, while the chronic phase included young adult (7 WT and 6 *Hivep3*-KO rats), adult (6 WT and 2 *Hivep3*-KO rats), and mature (5 WT and 7 *Hivep3*-KO rats). Statistical tests were performed using two-way ANOVA, following the model: value ∼ Genotype + Timepoint + Genotype x Timepoint.

### Flow cytometry phenotyping of *Hivep3*-KO rat

Before surface staining, 5×10⁵ cells per sample were incubated with diluted fixable viability dye (eFluor 506 or eFluor 450; eBioscience) for 20 min at 4 °C in the dark. After washing, cells were surface-stained with 30 μL of diluted fluorophore-conjugated antibodies for 25 min at 4 °C in the dark. For intracellular staining, cells were fixed and permeabilized for 45 min using the Foxp3/Transcription Factor Staining Buffer Set (eBioscience), followed by incubation with intracellular antibodies in permeabilisation buffer for 1 h. Fluorescence was measured on a BD FACSVerse flow cytometer, and data were analysed using FlowJo software. Statistical analyses were performed using two-way ANOVA, two-tailed t-tests, multiple t-tests, and the Mann–Whitney test; significance was set at p < 0.05.

The following rat immune cell populations were defined (markers shown in parentheses): T cells (CD45⁺TCRαβ⁺CD45RA⁻); CD4⁺ T cells (CD45⁺TCRαβ⁺CD4⁺); CD8⁺ T cells (CD45⁺TCRαβ⁺CD8⁺); B cells (CD45⁺TCR⁻CD45RA⁺); NK cells (CD45⁺TCR⁻CD161⁺); NKT cells (CD45⁺TCR⁺CD161⁺); MDSCs (CD45⁺TCR⁻CD161ˡᵒʷ); macrophages (CD45⁺CD11b⁺); granulocytes (CD45⁺RP1⁺); conventional dendritic cells (CD45⁺TCRαβ⁻CD103⁺CD4⁻ and TCRαβ⁻CD103⁺CD4⁺); plasmacytoid dendritic cells (pDCs; CD45⁺TCRαβ⁻CD45R⁺CD4⁺); effector T cells (CD45⁺TCRαβ⁺CD4⁺ or CD8⁺ with CD44⁺CD62L⁻); central memory T cells (CD45⁺TCRαβ⁺CD4⁺ or CD8⁺ with CD44⁺CD62L⁺); naïve T cells (CD45⁺TCRαβ⁺CD4⁺ or CD8⁺ with CD44⁻CD62L⁺); regulatory T cells (CD45⁺TCRαβ⁺CD4⁺Foxp3⁺). In addition, CD45⁺CD5⁺TCR⁻ (gate i), CD45⁺CD5⁺⁺TCR⁻ (gate ii), CD45⁺CD5⁺/⁺⁺TCRˡᵒʷ (gate iii) and CD45⁺CD5⁺/⁺⁺TCR⁺ (gate iv) subsets were further stratified into CD4⁺CD8⁻ (single-positive CD4), CD4⁺CD8⁺ (double-positive), CD4⁻CD8⁺ (single-positive CD8) and CD4⁻CD8⁻ (double-negative) populations.

### Tissue collection and histopathology

Multiple organ types were collected for histopathological examination: skin, thyroid, lymph nodes, colon, duodenum, ileum, kidney, liver, lung, and pancreas. Tissues were obtained from 9 rats aged over 1 year, comprising 6 *Hivep3*-KO animals (4 males and 2 females) and 3 WT animals (2 males and 1 female). All specimens were fixed in 10% formalin for 48 h and subsequently embedded in paraffin. At least two tissue sections (3 µm) per organ were stained with Haematoxylin–Eosin–Saffron (HES). All paraffin embedding, staining, and histopathological evaluations were performed by the APEX platform (APEX, INRAE, 2026, Plateforme d’Expertise en Anatomie Pathologique pour la Recherche).

### CD3 infiltration assay

3-µm tissue sections were baked at 60°C for 1 h and deparaffinized, and antigen retrieval was performed by incubating sections in citrate buffer (pH 6.0) at 96°C for 30 min. Sections were then blocked for 30 min with blocking solution (5% BSA and 5% normal goat serum diluted in PBS-Tween 0.05%) and incubated overnight at 4°C with anti-CD3 antibody (1:150, ab16669 Abcam) for immunofluorescence staining. After washing with PBS-Tween (0.05%), samples were incubated with secondary antibody (1:750, A11036 Invitrogen) for 1 h at RT and counterstained with DAPI for 10 min before mounting with an antifade mounting medium. CD3-positive cells were analysed and quantified using QuPath software. Statistical analyses were performed using the Mann–Whitney test; significance was set at p < 0.05.

### Quantification of serum cytokines IFNγ and IL-17A

Serum concentrations of IFNγ and IL-17A were quantified in 75 rat samples. Blood was drawn from 40 *Hivep3*-KO rats (20 males and 20 females) and 35 WT rats (22 males and 13 females) between 1 and 13 months of age. Following centrifugation, sera were separated, and cytokine levels were measured using the ProcartaPlex multiplex immunoassay kit (Cat. No. EPX010-30420-901, Thermo Fisher Scientific) on a Luminex platform at the CIMNA facility (Institut de Recherche en Santé de Nantes Université). The assay detection ranges were 5.58–22,220.6 pg/mL for IFNγ and 5.86–13,958.9 pg/mL for IL-17A. Serum samples were analysed either undiluted or diluted 1:2.

## Data and code availability

scATAC-seq and scRNA-seq data have been deposited (GEO XXXX). Published datasets used in this study include human scRNA-seq data from *Huisman et al. 2025*^16^, PolyKRT scRNA-seq from *Ragazzini et al. 2023*^17^, and rat scRNA-seq from *Stoljar et al., 2025* with HE-rat^37^.

## Supporting information

Table 1. Motif Enrichment analysis of human scATACseq TEC clusters

Table 2. Motif Enrichment analysis of mouse scATACseq TEC clusters

Table 3. Full list of DE genes in rat mTECHigh and cTECs_WT versus Hivep3-KO

Table 4. Full list of DE genes in rat mTECII_HE versus Aire-KO

## ACKNOWLEDGEMENTS

We thank Dr. Arnauld Sergé from the Centre de Recherche en Cancérologie de Marseille (Aix-Marseille Université, Marseille, France) and Thibaut Larcher from the APEX platform (APEX, INRAE, 2026, Plateforme d’Expertise en Anatomie Pathologique pour la Recherche, Nantes, France) for histopathological analyses. We thank Dr. Severine Bezie from the Center for Research in Transplantation and Translational Immunology (UMR 1064, Nantes, France) for her guidance with T-cell experiments.

## Author contributions

These authors contributed equally: Marie Massoda, Séverine Menoret and Axel You

## Supplementary information

**Supplementary Table 1.**
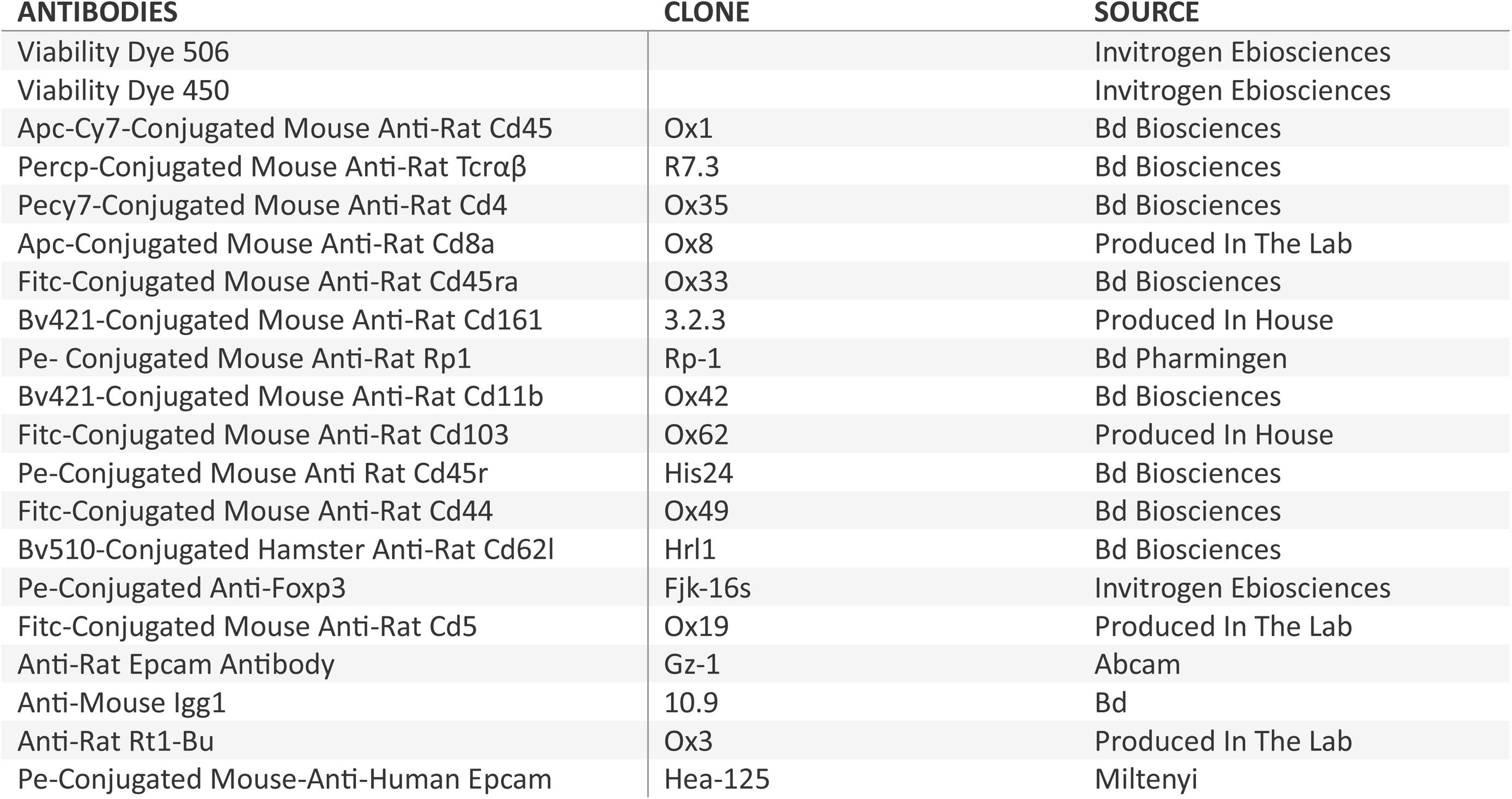
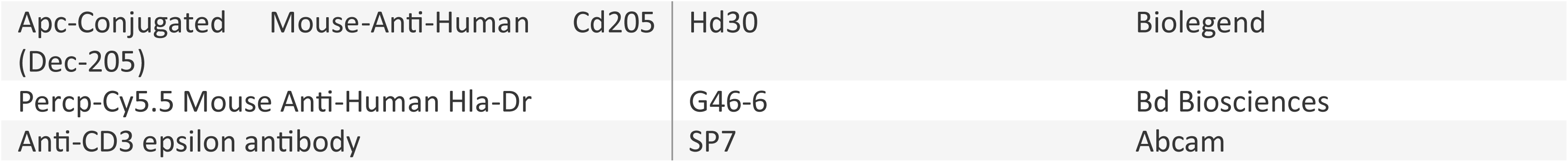
Antibodies used in the study.

---

**Table 1. Motif Enrichment analysis of human scATACseq TEC clusters**

**Table 2. Motif Enrichment analysis of mouse scATACseq TEC clusters**

**Table 3.Full list of DE genes in rat mTECHigh and cTECs_WT versus *Hivep3*-KO**

**Table 4.Full list of DE genes in rat mTECII_HE versus *Aire*-KO**

## Supplementary Figures

**Supplementary Figure 1.**
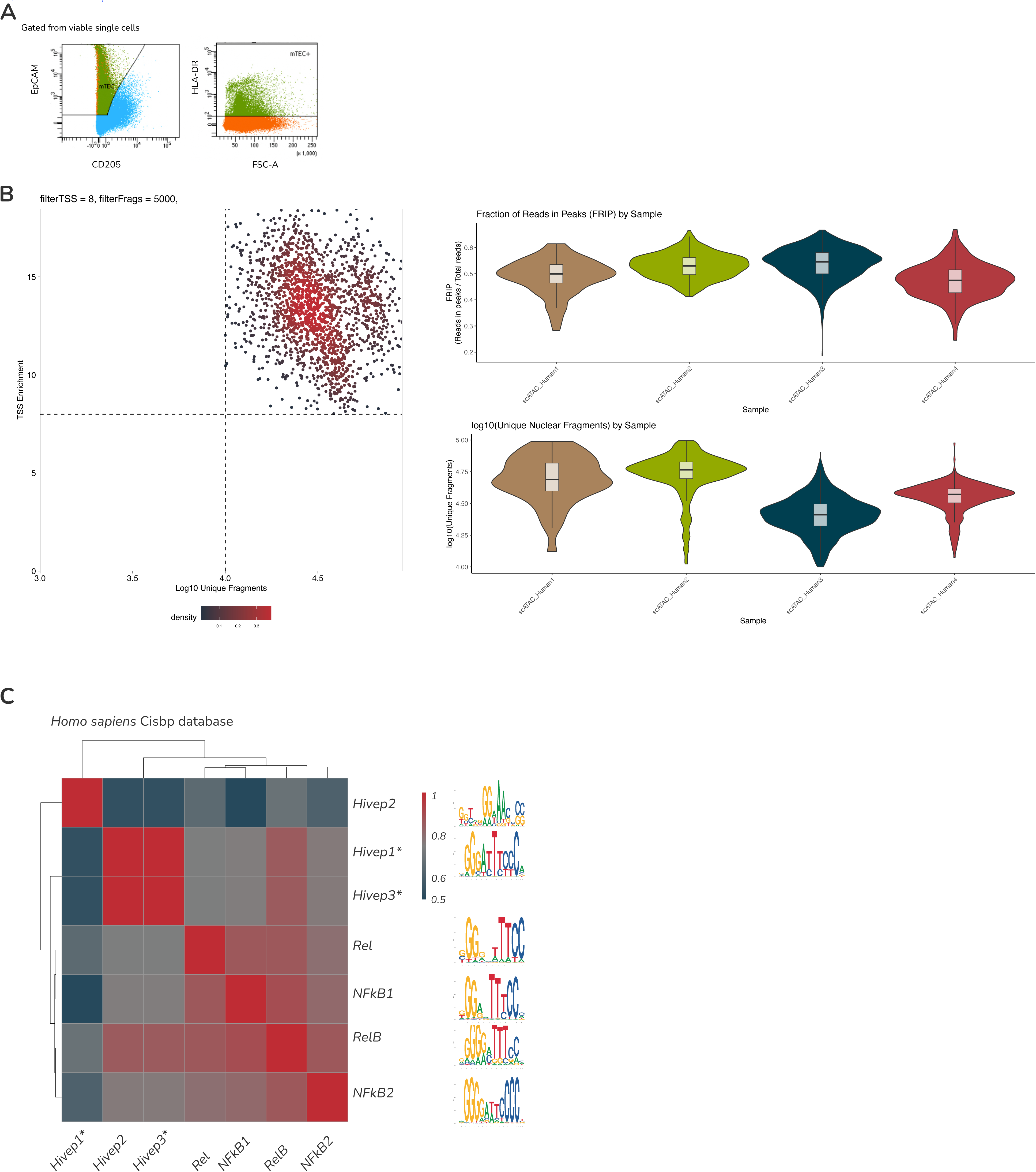
(A)Gating strategy used for the isolation of human thymic epithelial cell samples. Sequential gates were applied to exclude doublets and debris, select live cells, and enrich for epithelial cells (EpCAM⁺), followed by enrichment based on MHC-II expression. **(B)** Quality control metrics of scATAC-seq data from human TEC samples. Left: Dot plot showing Transcriptional Start Site (TSS) enrichment score (y-axis) versus log10(number of fragments) (x-axis) for every single nucleus. Right: Violin plots displaying fraction of reads in peaks (top) and log10(unique nuclear fragment) (bottom) across the human samples. The width of the violin indicates the density distribution. **(C)** Comparison of motif binding sites for NFκB (Rel, NFKB1, RelB, NFKB2) and HIVEP (HIVEP1/2/3). Data derived with ArchR from CIS-BP.

**Supplementary Figure 2.**
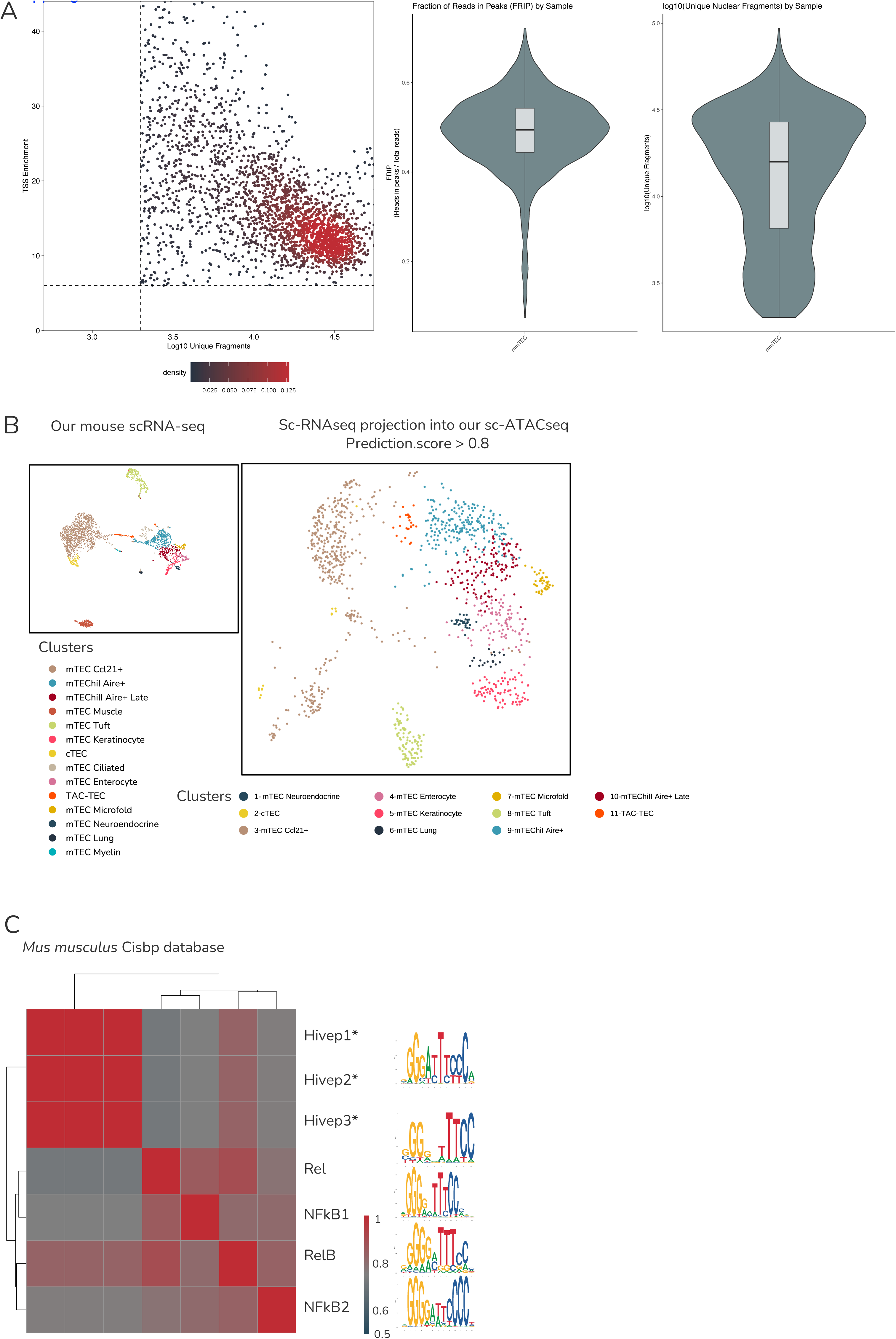
(A)Quality control metrics of scATAC-seq data from the mouse TEC sample. Left: Dot plot showing Transcriptional Start Site (TSS) enrichment score (y-axis) versus log10(number of fragments) (x-axis) for every single nucleus. Right: Violin plots displaying fraction of reads in peaks and log10(unique nuclear fragment) across the mouse samples. The width of the violin indicates the density distribution. **(B)** Integration of our scRNA-seq data from the mouse into our scATAC-seq dataset. On the left, UMAP of scRNA-seq. Each dot is a single cell. On the right, UMAP of the scATAC-seq with only the nuclei presenting a prediction score higher than 0.8. The predicted clusters are annotated and colour-coded. **(C)** Comparison of motif binding sites for NFκB (Rel, NFKB1, RelB, NFKB2) and HIVEP (HIVEP1/2/3). Data derived with ArchR from CIS-BP.

**Supplementary Figure 3.**
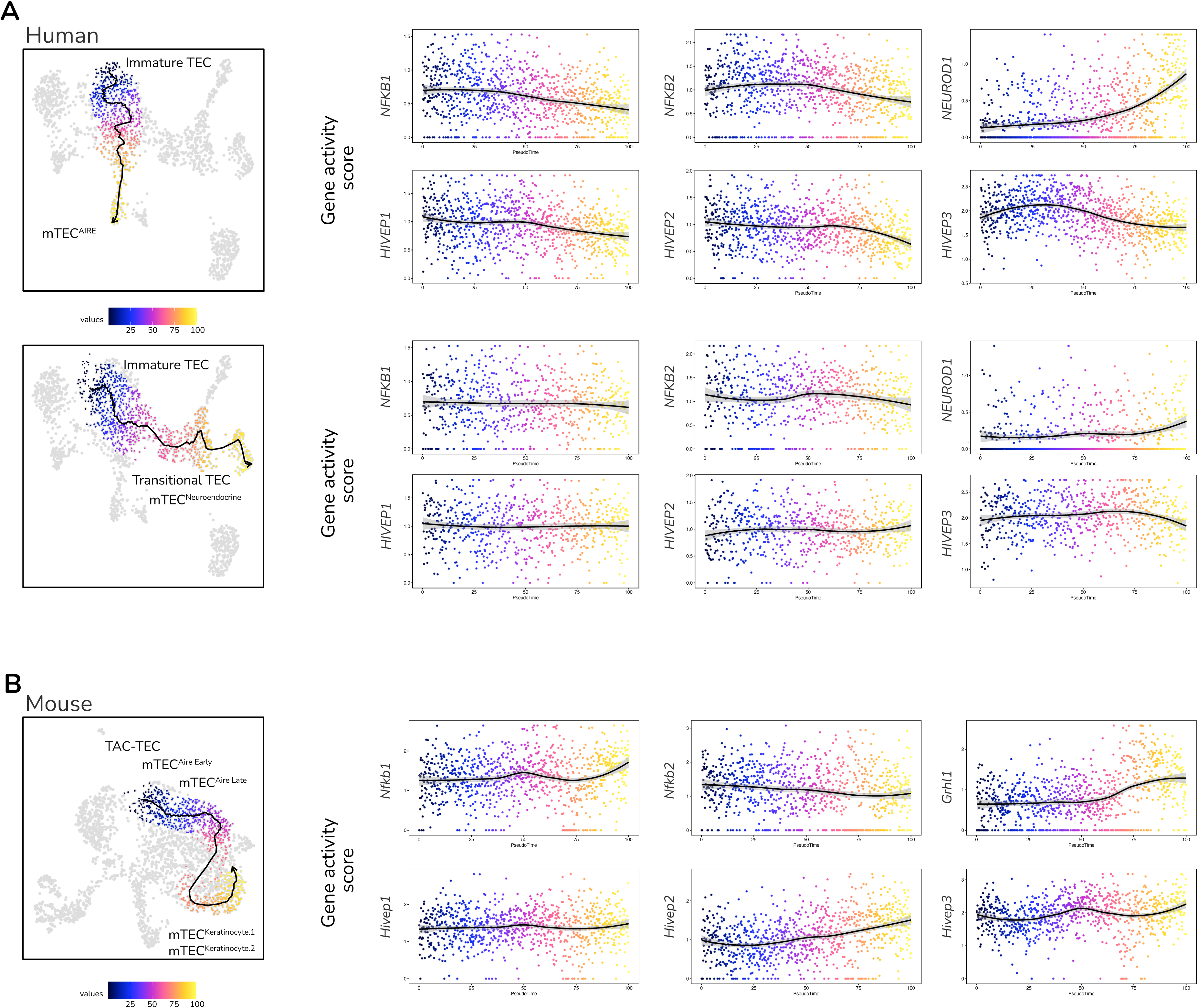
(A)Trajectory analysis in human scATAC-seq data spanning immature and mTEC^AIRE^ (top) or from immature and transitional mTEC to mTEC^Neuroendocrine^ for *Nfkb1/2*, *Hivep1/2/3* and *Neurod1* genes. **(B)** Trajectory analysis in mouse scATAC-seq data spanning TAC-TEC, mTEC^AIRE^ ^Early^, mTEC^AIRE^ ^Late^, mTEC^Keratinocyte.1^ and mTEC^Keratinocyte.2^ with individual feature plots for *Nfkb1/2*, *Hivep1/2/3* and *Grhl1* genes. Dot plots show the gene activity score for the corresponding transcription factors.

**Supplementary Figure 4.**
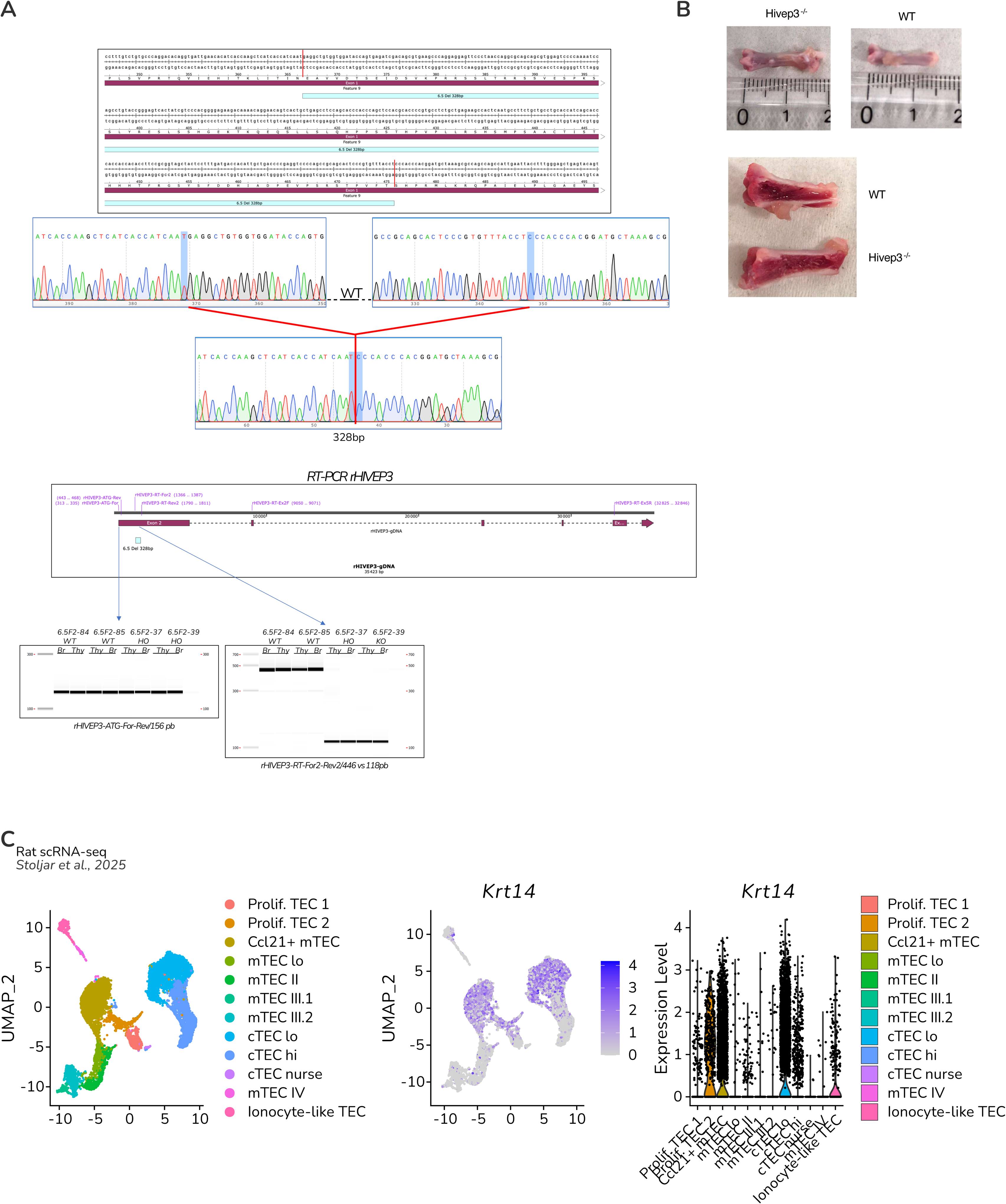
(A) Genomic representation of the localisation of *Hivep3* deletion in *Hivep3*-KO rats (top). RT-PCR on different regions of the *Hivep3* gene in WT and *Hivep3*-KO from the thymus (Thy) or brain (Br). **(B)** Representative images of whole (top) or bisected (bottom) femurs from *Hivep3*-KO and WT rats. **(C)** Uniform manifold approximation and projection (UMAP) of scRNA-seq of WT rat (*Stoljar et al., 2025*). Krt14 expression across the TEC population is represented in the feature plot (middle) or violin plot (right). Each dot represents a cell.

**Supplementary Figure 5.**
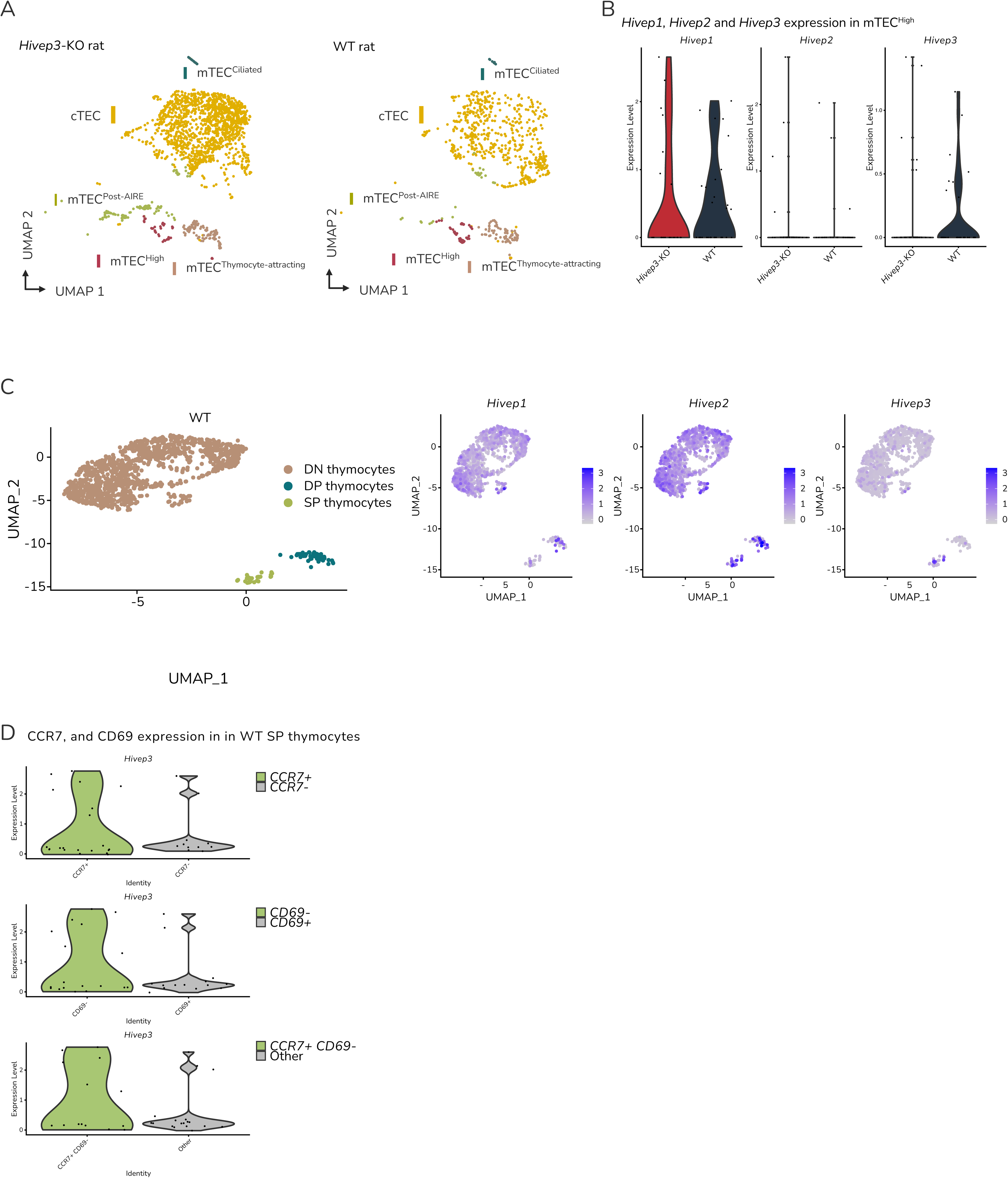
(A) Uniform manifold approximation and projection (UMAP) of scRNA-seq from *Hivep3*-KO (left) and WT (right) rats. Each dot represents a cell. **(B)** Violin plot of the expression of *Hivep1*, *Hivep2* and *Hivep3* in rat mTEC^High^ subsets in *Hivep3*-KO and WT. **(C)** Uniform manifold approximation and projection (UMAP) of scRNA-seq of WT rat thymocytes isolated from the thymus (left). Feature plot of *Hivep1, Hivep2*, and *Hivep3* across double negative (DN), double positive (DP) and single-positive thymocytes. **(D)** Violin plot representing *Hivep3* transcript level in Ccr7+, Cd69+ or Ccr7+ Cd69-single-positive (SP) thymocytes. Each dot represents a cell.

**Supplementary Figure 6.**
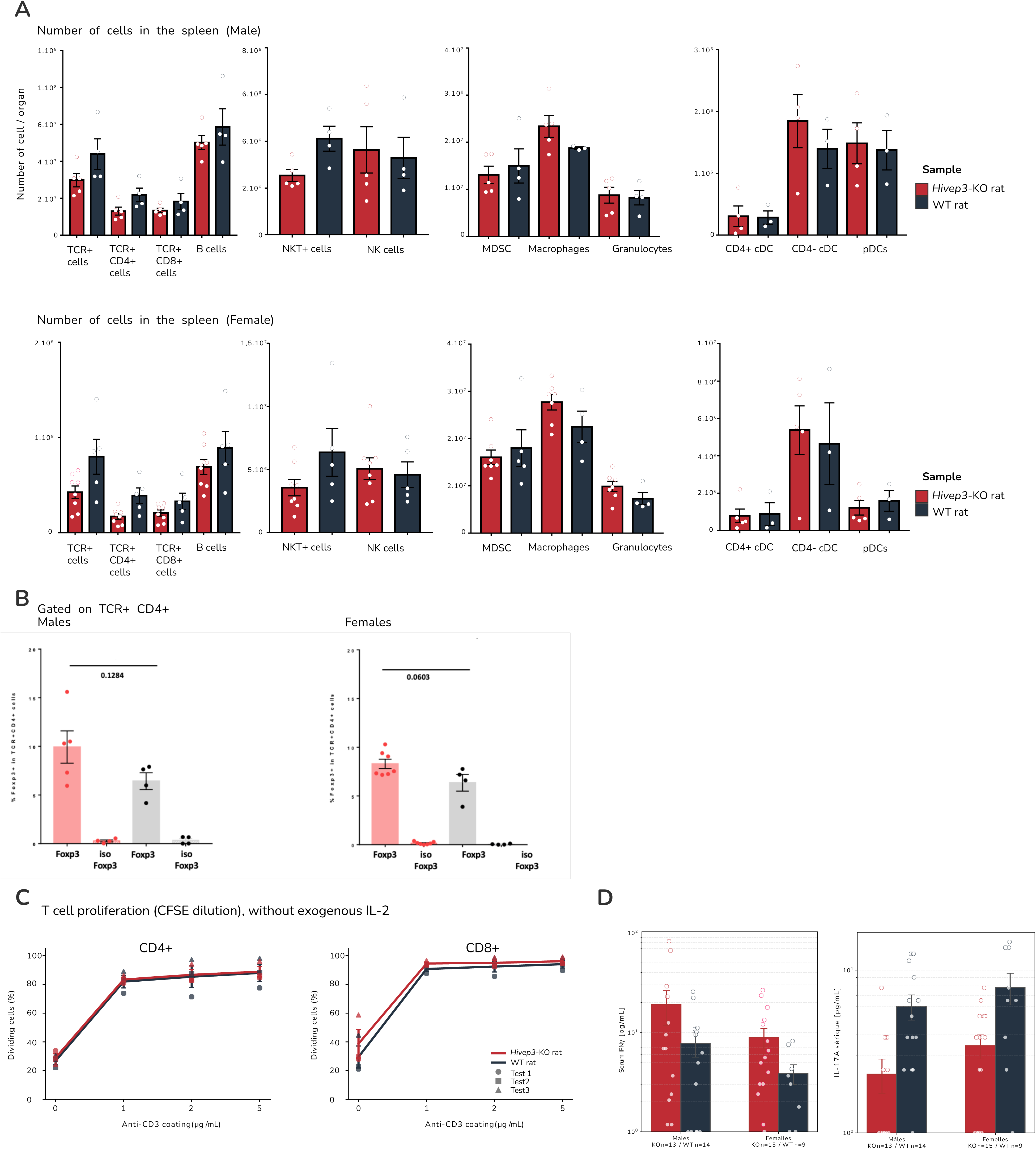
(A) Bar plot showing the number of splenic immune cells in male (top) and female (bottom) *Hivep3*-KO (red) and WT (blue) rats. **(B)** Percentage of regulatory CD4⁺ T cells in male (left) and female (right) *Hivep3*-KO (red) and WT (blue) animals. Each dot represents an individual animal. **(C)** CD3-induced dose-dependent proliferation of CD4+ (left) and CD8+ (right) lymphocytes. (n = 3). **(D)** Longitudinal analysis of IFNγ (left) and IL-17A (right) concentration (pg·mL⁻¹) in the serum of male and female *Hivep3*-KO (red) and control (blue) animals.

